# Cellxgene VIP unleashes full power of interactive visualization and integrative analysis of scRNA-seq, spatial transcriptomics, and multiome data

**DOI:** 10.1101/2020.08.28.270652

**Authors:** Kejie Li, Zhengyu Ouyang, Yirui Chen, Jacob Gagnon, Dongdong Lin, Michael Mingueneau, Will Chen, David Sexton, Baohong Zhang

## Abstract

To meet the growing demands from scientists to effectively extract deep insights from single cell RNA sequencing, spatial transcriptomics, and emerging multiome datasets, we developed cellxgene VIP (Visualization In Plugin), a frontend interactive visualization plugin of cellxgene framework, which greatly expanded capabilities of the base tool in the following aspects. First, it generates a comprehensive set of over eighteen commonly used quality control and analytical plots in high resolution with highly customizable settings in real time. Second, it provides more advanced analytical functions to gain insights on cellular compositions and deep biology, such as marker gene identification, differential gene expression analysis, and gene set enrichment analysis. Third, it empowers advanced users to perform analysis in a Jupyter Notebook like environment, dubbed Command Line Interface (CLI) by programming in Python and/or R directly without limiting themselves to functional modules available via graphical user interface (GUI). Finally, it pioneers methods to visualize multi-modal data, such as spatial transcriptomics embedding aligned with histological image on one slice or multiple slices in a grid format, and the latest 10x Genomic Multiome dataset where both DNA accessibility and gene expression in the same cells are measured, under the same framework in an integrative way to fully leverage the functionalities mentioned above. Taken together, the open-source tool makes large scale single cell data visualization and analysis more accessible to biologists in a user-friendly manner and fosters computational reproducibility by simplifying data and code reuse through the CLI. Going forward, it has the potential to become a crowdsourcing ecosystem for the scientific community to contribute even more modules to the Swiss Army knife of single cell data exploration tools.

## Introduction

Since the first single-cell RNA sequencing (scRNA-seq) study was debuted in 2009^1^, over 1600 scRNA-seq studies have been published to date, of which at least 72 studies have reported profiles in 200k or more cells^2^. The largest scRNA-seq study generated gene expression data of 4 million human cells^3^. It is foreseen that there will be a trend of increasing cell size to 500k or more in scRNA-seq studies. The sheer amount of data had brought challenges in visualizing and exploring such big data set interactively for scientists, even computational biologists.

Cellxgene^4^ is a leading open source scRNA-seq data visualization tool recommended in a recent evaluation^5^, which scales well in millions of cells and scores highly in user experience by leveraging modern web techniques with interactive features. Used by a large community of biologists, cellxgene proves itself in meeting the need of handling data interactively at scale. Except it lacks some essential plotting and analysis functions seen in scRNA-seq related command-line tools and publications that biologists are used to, hindering its utility and limiting scientists from taking advantage of ever accumulating scRNA-seq data sets to their full potential.

While bringing a more-fine grained assessment of each cell’s transcriptome, scRNA-seq loses spatial information during the cell dissociation process from a tissue. To gain deeper understanding of tissue cellular composition and cell-cell communication simultaneously, spatial transcriptomics methods^6^, e.g.: smFish^7^, MERFISH^8^, Spatial Transcriptomics^9^, seqFISH+^10^, NanoString GeoMx^11^, Slide-seq^12^, and 10X Visium^9^ are developed to allow the preservation of organization of tissue context and captures potentially cross talk between a variety of cell types within tissue^13, 14^. As the data grows in both volume and complexity involving histological imaging data and near single cell level transcriptomic measurements, presenting it in a digestible form for biologists is a daunting and challenging task. Currently, free or open-source spatial transcriptomics data visualization tools are either developed for computational experts comfortable of writing code in R or Python (Squidpy^15^, Giotto^16^, Seurat 4.0^17, 18^, and STUtility^19^) to interact with data, or offered as client end software (Loupe Browser from 10x Genomics) that requires installation on a personal computer that has very limited computing power. Although Loupe Browser and Giotto have user-friendly interactive features to overlay an image of stained tissue and gene expression on spatially distributed spots to explore multiple layers of information, none of these packages supports visualizing more than one samples at a time, which is needed for comparing replicates or selecting transcriptionally similar spots from multiple samples for downstream analysis, such as differential expressed gene (DEG) and spatial variable gene (SVG) analysis. Moreover, even for computational biologists, integrating spatial transcriptomics data with other analytical tools seamlessly poses a great challenge due to the limited capabilities of these tools.

Recently, 10X Genomics introduced multiome technology to simultaneously profile gene expression and open chromatin from the same cell to leverage gene expression markers to aid interpretation of epigenetic profiles, combine discovery of regulatory elements with gene expression to explore gene regulatory interactions, and obtain transcriptome and open chromatin information simultaneously to save samples and effort. Signac^20^ from the Seurat^17, 18^ family is the only tool we found to visualize this type of multi-modal data in a meaningful way by jointly plotting gene expression of genes and nearby peaks representing chromatin openness. However, bioinformatics skill is needed to master this command line tool.

To fill these gaps and to empower scientists to investigate these types of big data more efficiently, we developed a plugin of cellxgene named Visualization in Plugin, in short VIP to address the urgent needs of such essential functions for interactive visual exploration of scRNA-seq, emerging spatial transcriptomics and multiome data, and generation of publication-ready figures in a user-friendly way, especially for non-professional data analysts.

## Result

Cellxgene VIP’s strength lies in versatility, modularity, and scalability. Notably, it enhanced plotting functions significantly to generate violin, stacked violin, stacked bar, heatmap, volcano, embedding, dot, track, density, 2D density, Sankey and dual-gene plot in high-resolution by calling server-side plotting functions provided by Scanpy^21^, Seurat^17, 18^ and other general Python and R plotting libraries as illustrated in Figure 1 and supplementary tutorial. Innovative split plots in embedding (Figure 1a) and density plots (Figure 1g and Supplementary Fig. 13, see https://bit.ly/3j4SR0a) by annotations, such as cell types, make it easier to see which genes are differentially expressed in which cells. With the goal of becoming a platform to bridge users and various tools, cellxgene VIP not only leverages Python analytical capabilities, but also opens to another popular language R, which is demonstrated by a volcano plot (Figure 4b). In addition, interactive Sankey diagram (Figure 1f) and stacked barplot (Figure 1c) based on yet another language, JavaScript, enable investigation of the relationship between metadata and gene expression more intuitively.

**Figure 1.**
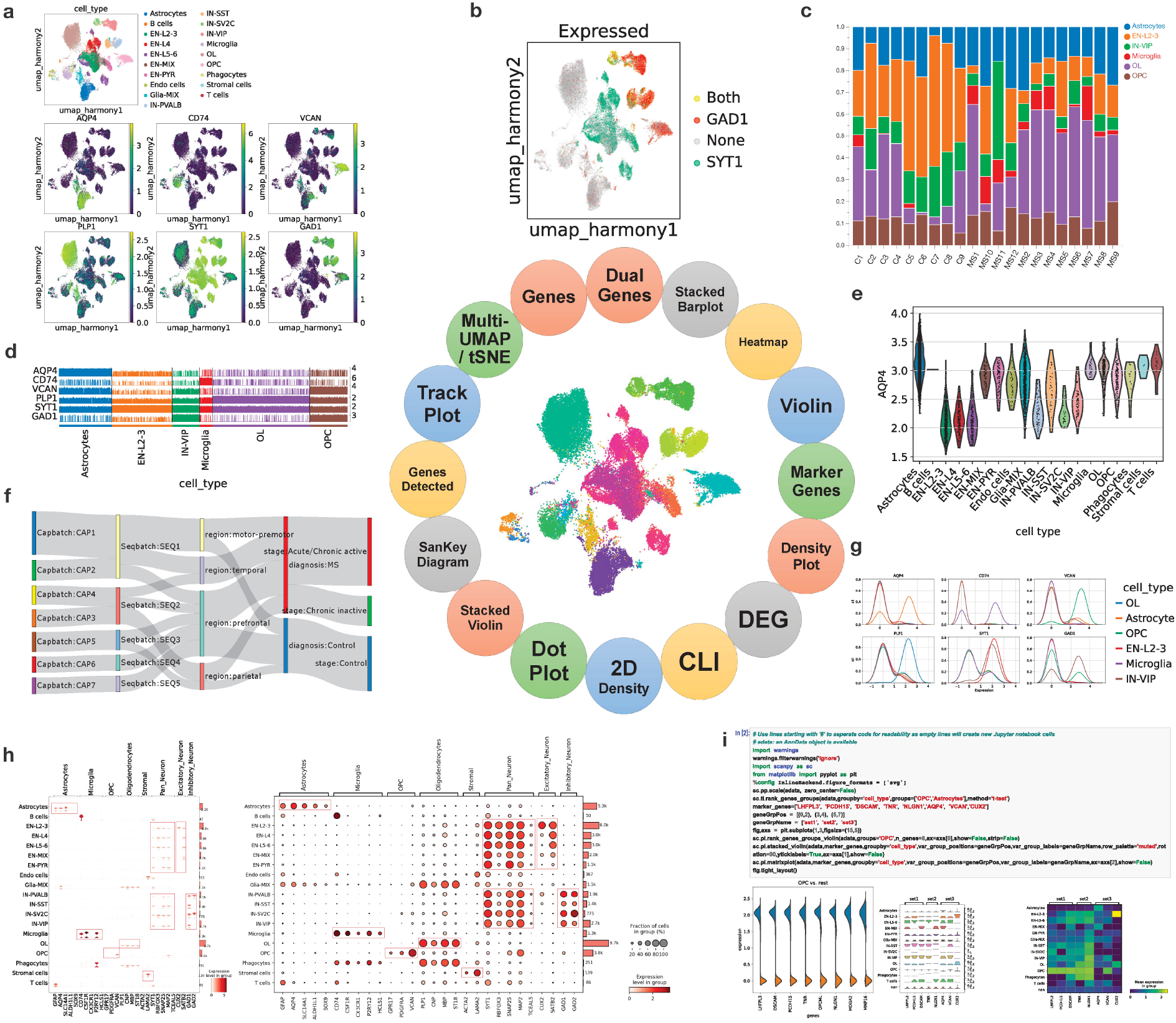
cellxgene VIP serves as an ecosystem of analytical modules that provide essential functions for interactive visualization and generation of publication-ready plots. Individual plots were assembled by bioInfograph^22^ with zoomable feature available at https://bit.ly/2QqdMg3 that is best viewed by Chrome. (**a**) **Multi-tSNE/UMAP plot** visually highlights which cells express cell markers on selected embedding, e.g., UMAP based on harmony batch correction in this example. (**b**) **Dual-gene plot** highlights cells expressing SYT1 and GAD1 (green SYT1 only, red GAD1 only, yellow co-expression of STY1 and GAD1), expression cutoff 2.2. (**c**) **Stacked barplot** demonstrates the fraction of each major cell type across each sample (C are Control and MS are MS patients). (**d**) **Trackplot** shows expression of lineage marker genes across individual cells in annotated clusters. (**e**) **Violin plot** shows the AQP4 gene expression across cell types. (**f**) **Sankey diagram** (a.k.a. Riverplot) provides quick and easy way to explore the inter-dependent relationship of variables in the MS snRNA-seq dataset^23^. (**g**) **Density plots** shows expression of marker genes across annotated clusters and split across cell types. (**h**) **Stacked violin** and **Dot plot** are the key visualizations of selected cell markers across cell types. They highlight their selective expression and validates the scRNA-seq approach and visualization method. (**i**) **CLI** exposed by mini Jupyter Notebook to provide maximal flexibility of performing various analytics on the whole or sliced single cell dataset.

To emphasize simplicity and flexibility, we paid great attention to user-friendliness by crafting the plugin to allow users to customize analysis and plotting parameters, set figure options like font size, image format and resolution, and pick a color palette for embedding as shown in Figure 2. To further strengthen the framework’s usability, the following convenient features are implemented such as displaying numbers of a brushed range, increasing the width of the central canvas to enlarge the working space, showing the description of the data set on the main window, and initializing the look-n-feel by a remote session file. Often, it is a challenge if not impossible to select interested cells by the cellxgene lasso tool in some situations, such as in ‘Spatial Transcriptomics’ where cells in a cluster could be separated in the space embedding, here the user can enable the cumulative selections in the global setting to select spots segment by segment. In another case, the user can turn on the scaling option to transform data to unit variance per gene (not zero mean due to the efficiency to handle sparse matrix) by calculating z-scores of the expression data using Scanpy’s scale function when it is appropriate. Further options of including genes with no variance and/or clipping at a threshold are provided for data scaling.

**Figure 2.**
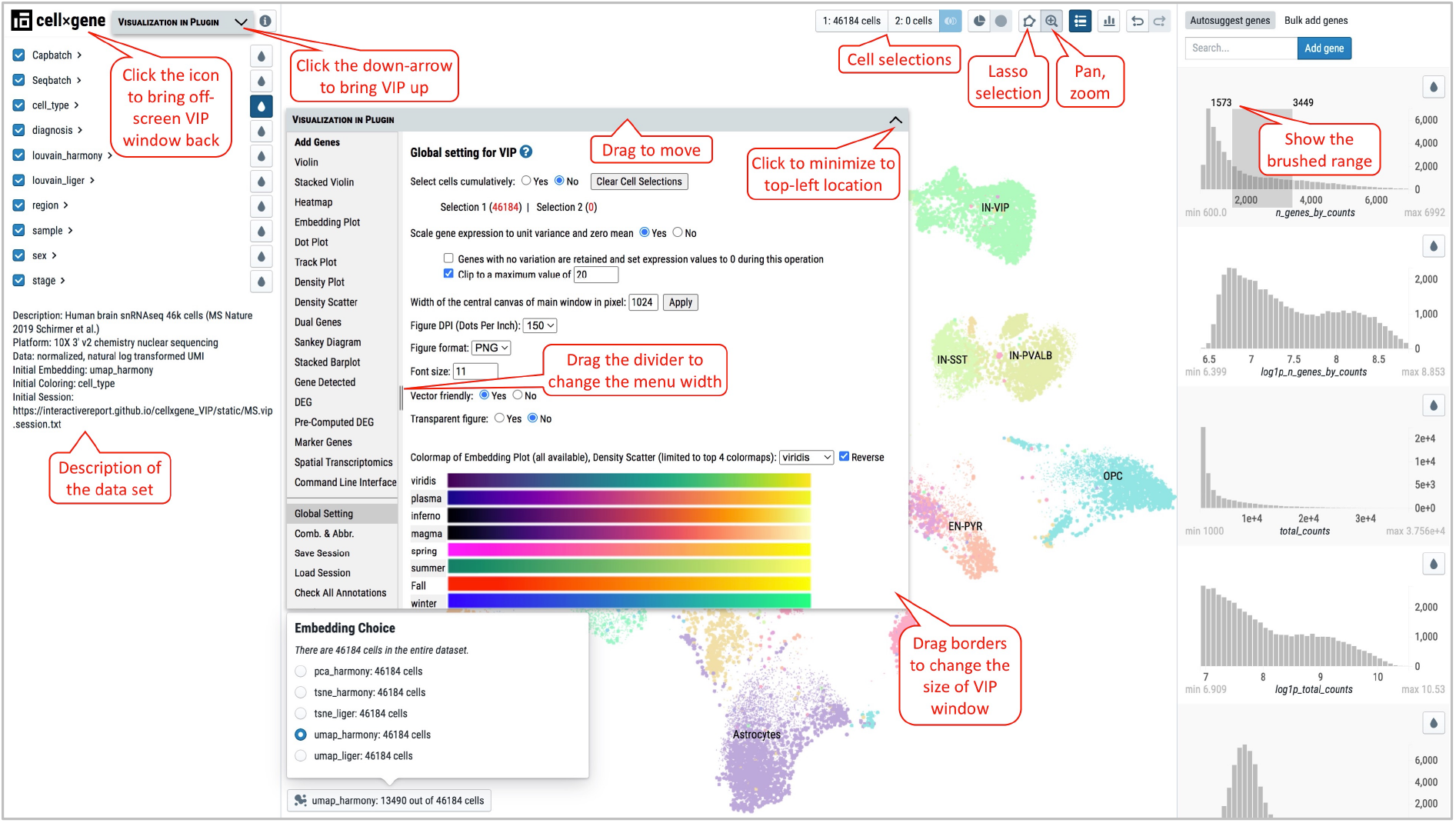
Main Window, VIP Window and setting parameters for figure plotting in Global Setting. Notable operations and features are highlighted in red call-out boxes. VIP windows can be fired up, minimized, moved, resized, and brought into scene by corresponding mouse actions.

To help biologists visualize and analyze 10X Visium Spatial Transcriptomics data, especially multiple samples/slices dataset, we developed a module in VIP to overlay the spatial spots on top of the Hematoxylin and Eosin (H&E) stained image (Figure 3a) and display gene expression of one or multiple genes on the spatial coordinates while other existing single cell visualizations are only applicable to spatial coordinate data. In VIP, the transparency of the H&E image or spatial spots can be adjusted to improve the visualization of the histology image, to zoom into specific regions of interest to allow better molecular insights under spatial context, to view derived layers/classification of glass slide spots with tSNE/UMAP, to allow manual cumulative selection of spots (Figure 3b) within regions of interest as groups or clusters by lasso tool, and to find differentially expressed genes between 2 regions occupied by selected spots. Besides overlaid visualization, viewing embedding image and gene expression profile side-by-side (Figure 3c) allows scientists to discover subtle histological features that usually are hidden in an overlaid setting. In addition, users can design the layout of sample slides to be merged by interactively changing grid location, rotation, and flipping of individual slides (Figure 3d) and then save the newly customized layout file for merging h5ad files by a Python script detailed in Method section.

**Figure 3.**
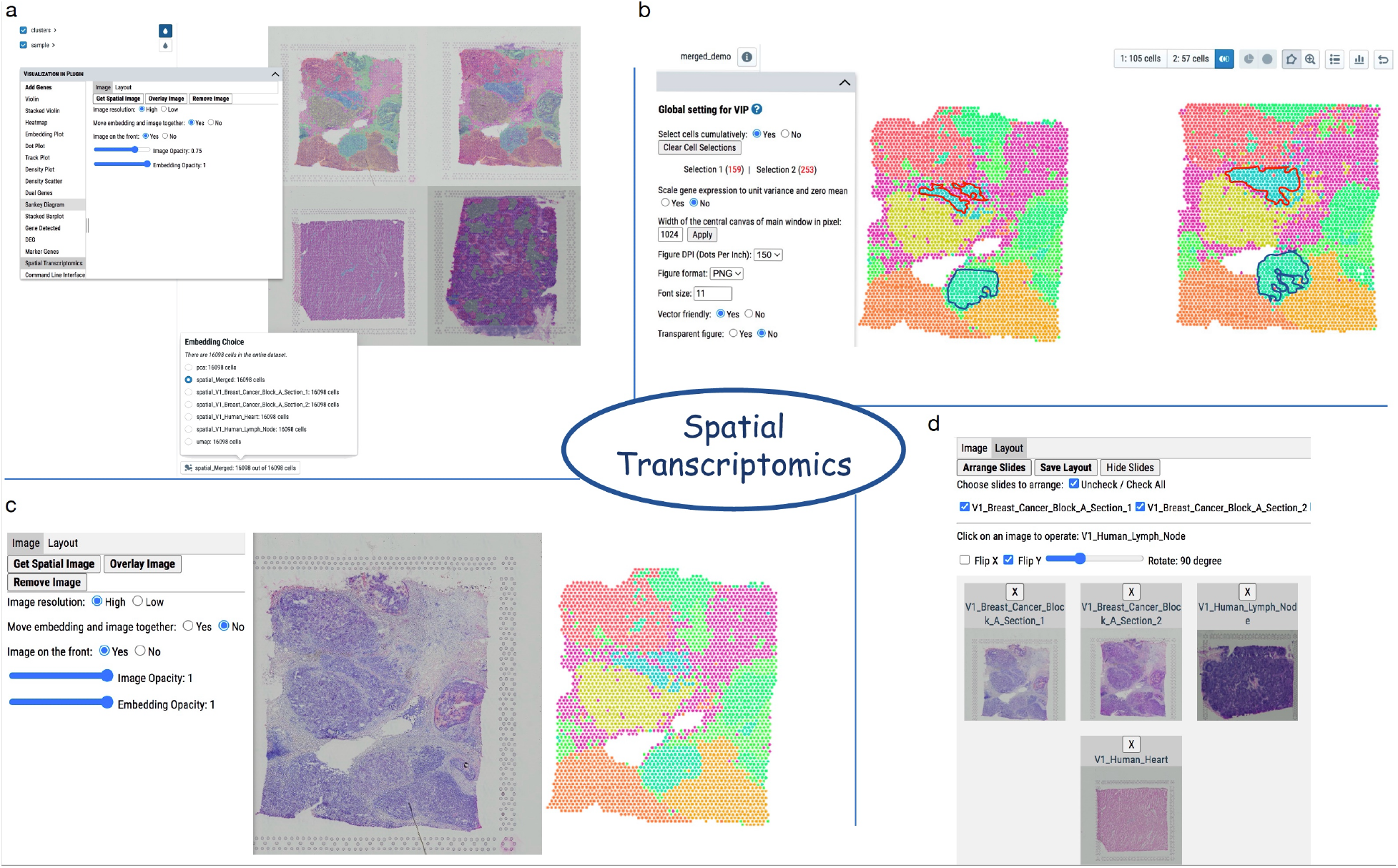
Advanced features of visualizing and processing of spatial transcriptomics data accessed by “Spatial transcriptomics” menu. (a) Overlapped view of merged spatial embeddings colored by clusters and histological images of multiple samples. The image can be loaded or removed by clicking on “Get Spatial Image” or “Remove Image” button, respectively. Overlapping order, opacity of embedding and image can be adjusted to obtain the optimal view. (b) Cumulative selection option is chosen in “Global Setting” to allow selections of cells from non-adjacent regions or multiple samples as shown by the red and blue irregular shapes drawn by lasso selection tool. Please note that numbers of cells in the main window refer to the current selection while numbers of cells in VIP window denote accumulated selections by multiple lasso operations. Downstream analysis such as differential gene expression can be performed between the accumulated cell groups. (c) Embedding and image could be reviewed separately by selecting “No” to “Move embedding and image together”. This way, subtle histological features obstructed in overlaid setting can be seen easily. (d) “Layout” menu provides a flexible way to re-arrange a slide by moving, rotating, flipping on X or Y coordinates as the original setting of the slide might not be the desired one, e.g., the third slide on the top row is rotated by 90 degrees and flipped on Y. A slide is activated by clicking on it or removed from the layout by clicking “X” on the top. In the end, the layout can be saved and used to create the custom merged h5ad file based on the on-screen design.

Beyond the visualization functions, we integrated diffxpy^24^ to provide more analytical capabilities in VIP. In addition to simple t-test and returning only a limited number of differentially expressed (DE) genes in cellxgene’s DE analysis, users can get the full list of DE genes by calling any of the methods implemented in diffxpy, namely Welch’s t-test, Wilcoxon rank-sum test, and Wald’s test. Volcano plots in both interactive and static forms with fold change and p-value are also generated to enable global view of biological perturbation (Figure 4b). Gene Set Enrichment Analysis (GSEA) can be elected to run for the user in DE analysis as well. Since the response speed is one of the critical concerns of the web application, we used fgsea implementation^25^. Users can also download the pre-defined gene sets (MSigDB: symbol version) from Broad Institute or generate their own. Cellxgene VIP also allows the user to load and visualize pre-computed DE gene lists by any other single cell differential analysis algorithms requiring long computational run time, such as glmmTMB^26^ or NEBULA^27^.

**Figure 4.**
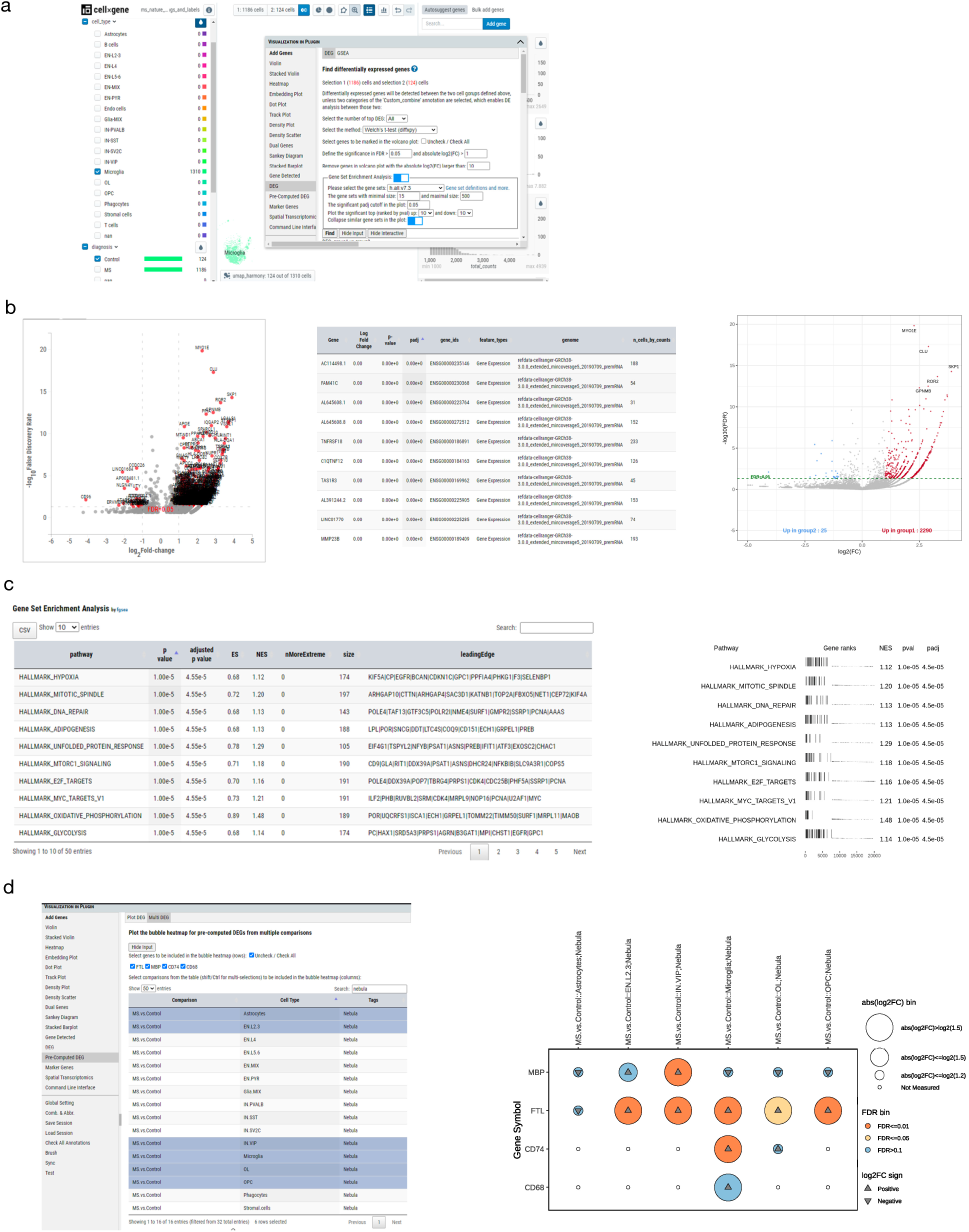
Differential Gene Expression analysis. (a) User could select microglia cells and define selection 1 as MS (1186 cells) and selection 2 as Control (124 cells). DEG analysis could be performed by using Welch’s t-test from diffxpy implementation. GSEA could be enabled too. All with default parameters except showing full DEG result and select hall mark gene set (h.all.v7.3) from GSEA. (b) Result of the simple DEG analysis. 1. Interactive volcano plot, 2. Searchable DEG table, 3. Static volcano plot. (c) GSEA results. (d) Bubble plot to explore known list of gene DE results. Here we would like to see how a list of genes “FTL, MBP, CD74 and CD68” DE results across a defined set of pre-computed comparisons. From the bubble plot, user can easily see FTL is statistically significantly differential expressed in Microglia, Layer 2 and 3 Excitatory Neurons (EN.L2.2), Oligodendrocytes (OL), OPC and VIP+ inhibitory neurons (IN.VIP) with fold change > 1.5 and FDR < 0.05. While CD74 is up regulated in Microglia and MBP is up regulated in VIP+ inhibitory neurons.

For demo purpose, Schirmer et al.^23^ data was re-processed from fastq files for cellxgene VIP visualization (details of the reprocessing in iPython notebook format could be found from our GitHub repository at https://bit.ly/2CeUHtO, while the demo site is available at https://cellxgenevipms.bxgenomics.com). In this study, Schirmer et al. acquired brain blocks from 12 MS and 9 control individuals and generated snRNA-seq data to study the disease mechanism of Multiple Sclerosis. From the example below (Figure 4), a user would be able to quickly look for trends by performing fast statistical tests online as shown in Figure 4a and then confirm by more sophisticated and time-consuming statistical tests ran offline (Figure 4b). To illustrate the importance of using a statistically rigorous test after the quick investigation, we use the finding of FTL (ferritin light chain) gene as an example. FTL is responsible for storing iron in a soluble, non-toxic, readily available form, which is important for iron homeostasis. Iron is taken up in the ferrous form and deposited as ferric hydroxides after oxidation. The OpenTargets platform^28^ shows a connection from FTL to Multiple Sclerosis (https://bit.ly/3Krq6qN). From MS disease pathology, accumulated phagocytosed iron in a brain lesion is associated with clinical deterioration^29, 30, 31, 32, 33, 34^. In this dataset, FTL gene shows weak upregulation in MS vs Control in microglia from t-test with log2 fold change of 0.49. Since a generalized linear model (glm) with “age”, “sex”, “mitochondrial fraction”, “capture batch” as covariates is more appropriate, NEBULA^35^ and glmmTMB^36^ methods were applied, and FTL is shown to be strongly upregulated in MS microglia cells with log2 fold change of 2.46. Further, gene set enrichment analysis (GSEA) is also available in VIP for understanding of biological pathway context (Figure 4c). Finally, a bubble plot visualization provides the user the ability to directly query a list of genes from the DE analysis results. It allows quick global overview of differential expression patterns for a list of genes across many comparisons from the pre-computed DE results as shown in Figure 4d.

**Figure 5.**
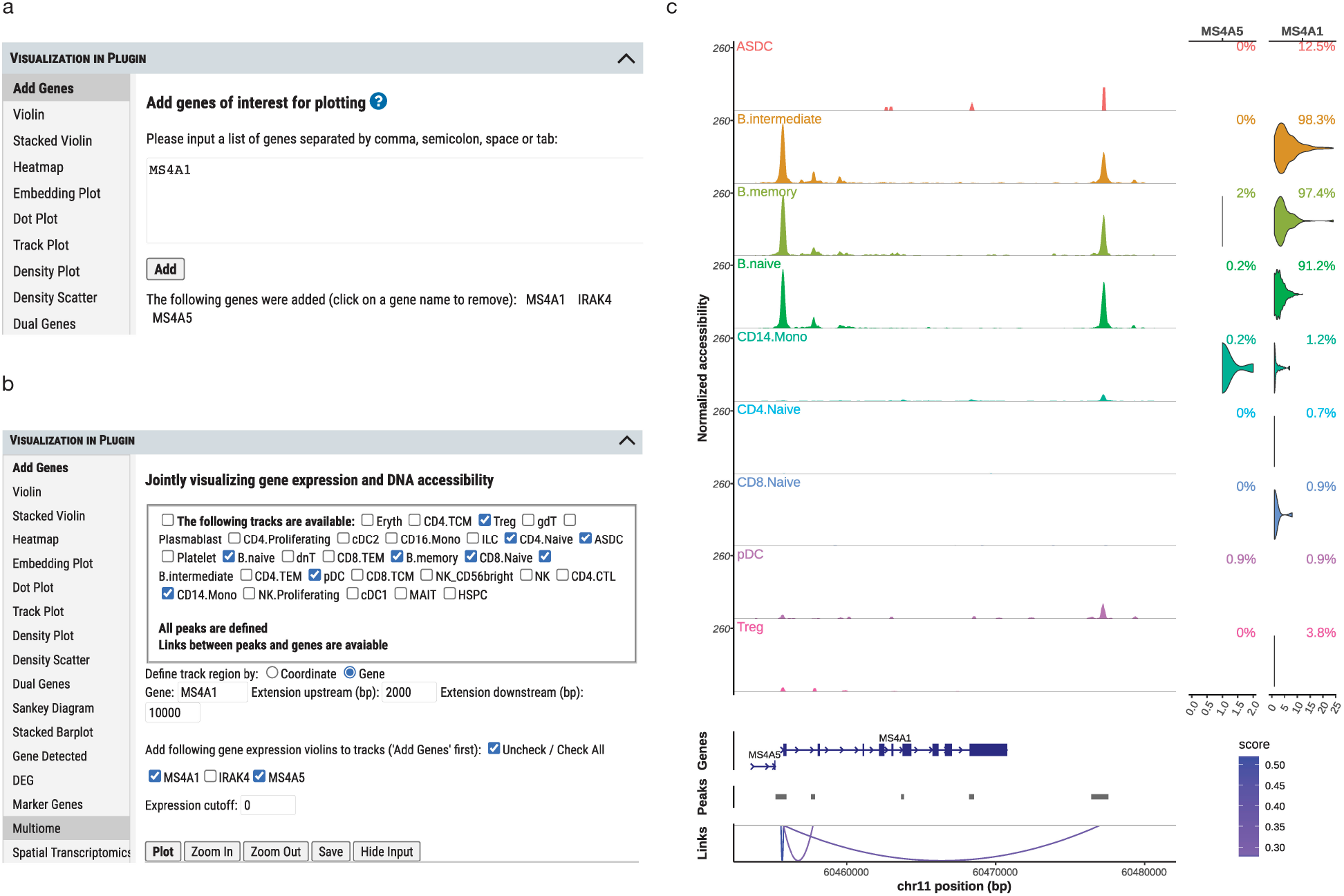
Visualization of multiome (scRNA-seq and scATAC-seq) data. (a) Add genes to be investigated first under “Add Genes” menu. (b) In “Multiome” module, user can select cell type specific tracks of DNA accessibility, regions of DNA, and added genes before “Plot” action. (c) Joint plot of genome browser track of DNA accessibility peaks from bigwig files and violin plot of gene expression. The percentage of cells expressing a gene above the cutoff in log scale is shown in the up-right corner of a violin plot. Pre-computed Pearson correlation between the expression of a gene and the accessibility of each peak is displayed as a curved line color-coded by the score scale in the legend.

scRNA-seq provides insight into transcriptomic heterogeneity and unveils unknown cell types in cellular populations^37, 38, 39^, while scATAC-seq dissects the transcriptional regulation network in mixed cell populations^40, 41, 42^. The integrative analysis of multimodal single-cell data including RNA transcription information and chromatin profiles provides an unprecedented opportunity to investigate the cell type-specific gene regulation and cell-to-cell variance in gene expression^43, 44, 45^. Using a cryopreserved human peripheral blood mononuclear cells (PBMCs) dataset^46^, we illustrated how cellxgene VIP allows users to explore this multiome dataset interactively (Figure 5). IGV styled genome browser visualization of combined DNA accessibility and gene expression data at one of the B cell marker genes, MS4A1 locus, shows the differential DNA accessibility among B cells and other cell types. The displayed link shows how DNA accessibility at the locus correlates with the expression of nearby genes within a given distance from the gene transcriptional start site, where peaks representing DNA chromatin openness in all B cell classes are highly correlated with the expression level of MS4A1 gene but not the other gene, MS4A5.

To further empower advanced users with programming skills, a Command Line Interface (CLI) is provided to offer unlimited analytical capabilities by directly utilizing any Python and/or R packages such as Seurat or any other R packages installed on the server (sample vignettes in CLI) through a mini Jupyter notebook environment as shown in Figure 6. All cells are used in vignette 1 (V1) while only T cells are selected in V2 and 3 for generating the result quickly. Computationally intensive and time-consuming jobs are not recommended to run in CLI. It is mainly to prototype and perform bioinformatics analysis in a small subset of cells or generate plots that is not provided by other interactive functional modules. Moreover, by sharing a collection of well-test vignettes with other analysts or even biologists, use cases of data reuse will be greatly expanded, e.g., general biologists could just change gene names in vignette 1 to generate a joint plot for genes of interest without seeking assistance form a computational biologist. Even for computational biologists, it is a great learning platform to share code snippets and workflows with each other.

**Figure 6.**
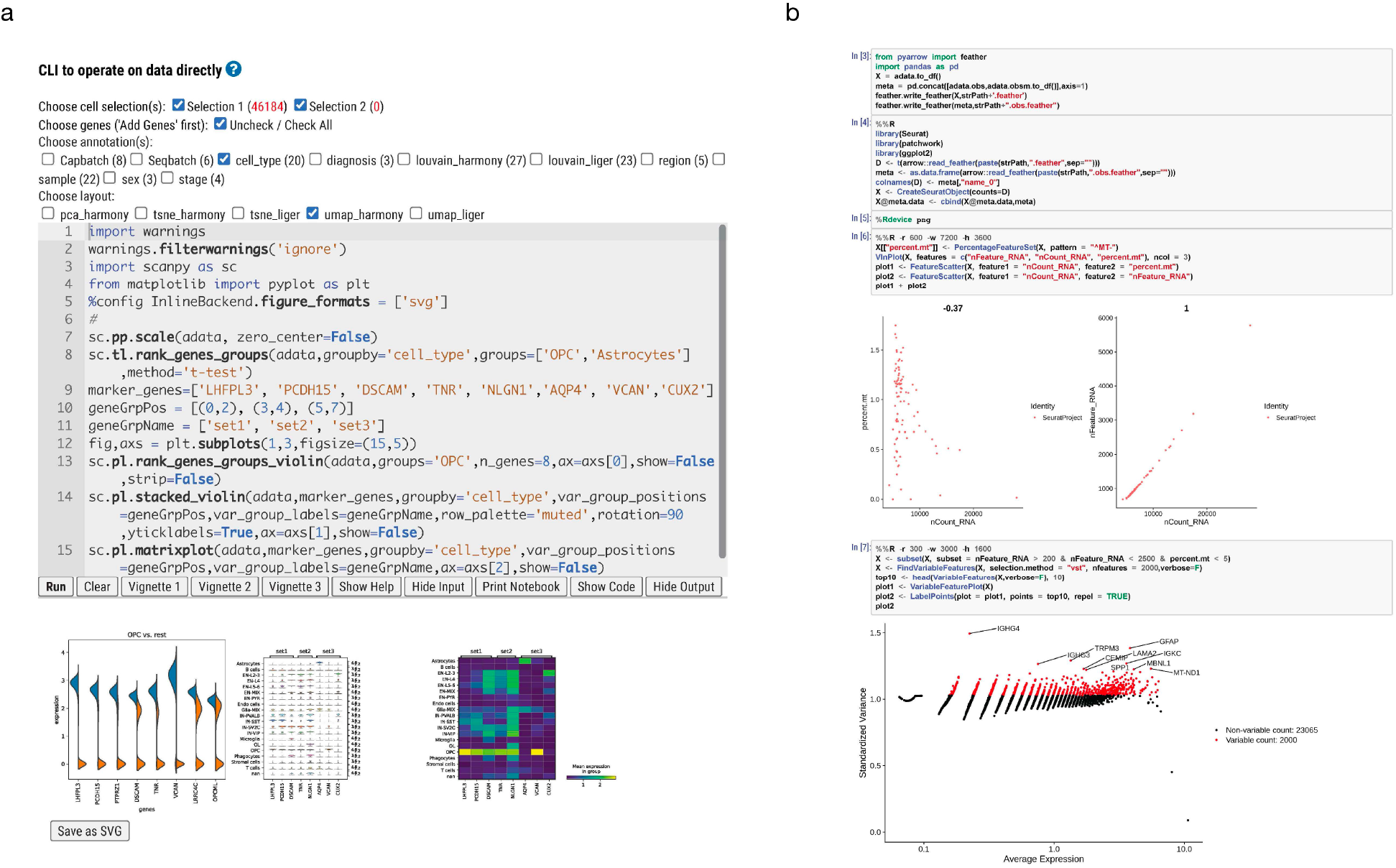
Command Line Interface (CLI) and demonstration of code examples by vignettes. (a) Pure Python code provided by clicking on “Vignette 1” operates on all cells by “Run” button to generate a joint plot in SVG format that can be readily saved. Through GUI, a user can select a limited number of annotations and layouts to minimize data trafficking between the server and the in-memory h5ad file. Other functional buttons are for additional convenience, such as “Hide input” will increase the space for viewing the result. (b) Partial output of “Vignette 2” that combines both Python and R code blocks in a Jupyter notebook like environment. Since the huge in-memory data needs to be converted to a file that R code will operate on, only a subset of cells is selected for the demo purpose. “Vignette 3” will generate a similar report in R markdown form that is not shown here.

Loading half-million-cell dataset even with sparse matrix compression applied would take up to approximately 20G of memory. Standard Mac/PC laptop would not be able to load such a dataset into memory, not to mention doing additional interactive exploration and plotting. Within cellxgene VIP, all heavy lifting functions such as differential gene expression analysis are executed on the server side that in turn greatly reduces the burden on the client browser side.

To facilitate reproducible research, sessions can be saved including all settings in the VIP window and cell selections in the main window, which can be loaded later to restore settings from previous analysis, with options to load from a URL or local session file and follow by reproducing plots in the previous session in one click (Supplementary Materials for more details).

Cellxgene VIP also initializes cellxgene embedding, coloring and brief description of the data (Supplementary Methods), sync cell selection, gene input from the main window to VIP automatically, save and load sessions, and create combinatorial annotations and abbreviations (Supplementary Fig. 24, https://bit.ly/376k28w). Further, we made cellxgene VIP open source and free of charge to facilitate broader scientific collaborations on large scale scRNA-seq data sets.

In summary, our web-based interactive tool is built upon cellxgene but greatly extended its plotting and analytical capabilities by integrating state-of-the-art tools in this field. It allows users with no programming skills to rapidly explore scRNA-seq, Visium spatial transcriptomics, and multiome data interactively and create high-resolution figures commonly seen in high-profile publications. Furthermore, it is the first tool to the author’s best knowledge to allow computational biologists to write their own code to communicate with the hosting server via a mini Jupyter notebook interface to perform advanced analysis or provide code snippets for others to reuse in the same environment. It stimulates unlimited capabilities even beyond the rich set of plotting and analytical functions already provided in the tool.

## Methods

### Software architecture

Cellxgene VIP uses a server-client architecture that enables users to take full advantage of an interactive web browser-based interface to set parameters for analysis, plot figures and program in CLI while the heavy lifting computational tasks are performed on the server side. On the server side, the most popular scRNA-seq and spatial transcriptomics data processing tools, Scanpy and Seurat, are integrated for data analysis and visualization. Various JavaScript libraries including D3, jQuery, Crossfilter, and Ace were used to provide the interactive user experience.

VIP makes minimal modifications of cellxgene source code. It takes advantage of the cached data by cellxgene server to avoid data reloading which is time consuming for a large data set in gigabytes. On the client-side, a non-invasive JavaScript panel (Supplementary Method 1, https://bit.ly/3rftBZA) was elegantly plugged in to bridge the communications between the user and server through the web browser store (Supplementary Method 3, https://bit.ly/37zXeOq). With extensibility in mind, we decoupled the development of the plugin and cellxgene as much as possible, so it is easy to incorporate additional tools in Python or any other languages like R on server side while taking full benefits of the interactive user experience offered by both cellxgene and the plugin. Located on the same web page, the VIP works seamlessly with cellxgene main window through real time data exchange by JavaScript, such as synchronizing cell selections and showing ranges of brushing on histograms of continuous annotations. With such design, developers can easily add new functions to the plugin without touching the cellxgene code base at all. In the end, we provide easy-to-use scripts to make necessary modifications mentioned above and install cellxgene VIP components, R packages and Python modules on the server side.

### Input data format

AnnData (https://anndata.readthedocs.io) format developed for Scanpy is used along with a text file with custom description and initial settings. We follow the same input requirements as cellxgene^4^. To start cellxgene, a h5ad file meeting the following AnnData specifications is required: 1. anndata.X is the raw/normalized expression matrix; 2. annadata.obsm contain at least one embedding (with prefix “X_”, such as “X_tSNE”); 3. Cell barcode or gene IDs must be unique. To provide more details about the data set and set initial options of visualization, we create an optional description file to add a study dataset description at the bottom of the categorical annotations on the left panel. The description file is required to be the same name as the h5ad file with ‘txt’ as extension within the same directory. The content of an example description file is as follows:

*Description: Human brain snRNA-seq 46k cells (MS Nature 2019 Schirmer et al.)*

*Platform: 10X 3′ v2 chemistry nuclear sequencing*

*Data: normalized, log transformed UMI*

*Initial Embedding: umap_harmony*

*Initial Coloring: cell_type*

*Initial Session: https://interactivereport.github.io/cellxgene_VIP/static/MS.vip.session.txt*

Colon is used as the delimiter for VIP to parse the initial settings for embedding and coloring. In our demo, the default layout is “umap_harmony” while cells will be colored based on “cell_type” categories. If keyword “scale” is not found in “Data” line, VIP will set the figure option “Scale data to unit variance for plotting” to “Yes”. The value of “Initial Session” will populate the “Load from a URL” input in “Load Session” interface.

### Sample data used for demonstration of cellxgene VIP functionality

Human MS brain snRNA-seq dataset raw data (fastq, sample metadata and cell type identification files) from Schirmer et al.^23^ was downloaded and re-processed with 10x cellranger 3.0.2 (count) by mapping to human pre-mRNA genome reference (GRCh38 reference - 3.1.0, July 24, 2019). Quality control criteria were applied as follows: filtered out cells that have less than 1000 counts, or more than 40000 counts, and kept cells with < 0.2 mitochondria gene counts. Harmony^47^ and Liger^48^ were used to correct for batch effects.

Louvain clustering method^49^ was used to cluster cells into different subgroups (clusters) on both batch corrected results. Both tSNE^50^ and UMAP^51^ coordinates are provided. Authors’ cell type annotation was in great concordance with the Louvain clustering result, except we decided to merge OL-A, OL-B and OL-C clusters into OL (Oligodendrocyte) and EN-L2-3-A and EN-L2-3-B clusters into EN-L2-3 (Excitatory Neuron Layer).

### Integrated gene set query

Cellxgene VIP equips a helper tool, xGenesets that provides an Application Programming Interface (API) for web-based applications to select a list of genes in a gene set. The gene sets are defined based on pathways, gene sets, and categories from various sources, including KEGG Pathways^52^, WikiPathways^53^, Small Molecule Pathway Database^54^, Reactome^55^, Gene Ontology^56^, Molecular Signatures Database^57^ from Gene Set Enrichment Analysis (GSEA^58^), and LIPID MAPS Proteome Database^59^. xGenesets currently has 175,537 gene sets stored in a MySQL database, covering gene sets for species including human (115,497 records), mouse (30,554 records) and rat (29,486 records). xGenesets features auto-completion for quick gene set selection, gene set database browsing, and the full member gene list of any gene set. It can be easily embedded into an online application by leveraging jQuery plugin, DataTables, and bootstrap (either v3 or v4). An example of using xGenesets API can be found at https://bxaf.net/genesets.xGenesets API is freely available for all usages.

### Generation of combo annotations

Celllxgene VIP also provides a fast annotation combination tool to label cells across selected categories. A new category, “Custom_combine”, will be automatically created to annotate cells across different categories, such as cell type and diagnosis. The user can also re-order the annotations (e.g., EN-L2-3:MS, EN-L2-3:Control) in this category which can be used in VIP plotting functions. Cells not belonging to any combined annotations will not be considered in plotting.

### Differential gene expression

Three statistical test methods of differential gene expression analysis are provided including Welch’s t-test (both cellxgene and diffxpy implementations), Wilcoxon rank test, and Wald’s test. The statistical test results are presented in a table format including log2 fold change, p-value, and multi-testing adjusted p-value by Bonferroni in cellxgene or Benjamini-Hochberg in diffxpy. Notably, we provide users with simple test methods owing to quick response time within the interactive framework. However, more advanced statistical methods developed for differential gene expression analysis of scRNA-seq data such as glmmTMB^26^ or NEBULA^27^ are recommended even if it takes much longer computational time to get results considering that there would be covariates that need to be modeled in a proper statistical test. To accommodate data exploration of offline results, we developed a loading script for users to load and explore pre-computed results interactively inside the VIP environment in table, interactive volcano, and bubble heatmap forms.

### glmmTMB differential analysis

As an example of an advanced testing method, we applied the glmmTMB algorithm^26^ to the data of Schirmer et al^23^. The glmmTMB method uses a generalized linear mixed model framework combined with the template model builder to determine differentially expressed genes. This method is very flexible allowing the user to adjust for covariate effects and model subject to subject or sample to sample variance components. The user can choose a variety of response families, dispersion models, zero inflation models, and variance/covariance structures. Unfortunately, this approach is very computationally demanding so its DEG results were pre-computed and then visualized using cellxgene VIP.

### NEBULA differential analysis

As a second example of an advanced testing method, we applied the NEBULA algorithm^27^ (both NEBULA-LN and NEBULA-HL) to the data of Schirmer et al^23^. The NEBULA method uses a generalized linear mixed model framework to determine differentially expressed genes. This method is flexible allowing the user to adjust for covariate effects, to account for subject-level overdispersions, and to account for cell-level overdispersions. A large sample approximation in NEBULA-LN and the h-likelihood method of NEBULA-HL allows NEBULA to be computationally efficient and to be much faster than other single-cell level DEG tools. After calculating the DEGs for the data of Schirmer et al^23^, the results were visualized using cellxgene VIP.

### Filtering criteria of DE analysis

For the NEBULA and glmmTMB differential analysis, the counts data of Schirmer et al^23^ was imported into R and two rounds of QC filtering were applied. In the first round, filtering was applied to the entire count matrix. We require: 1) a library size between 200 to 20M, 2) genes must be expressed in at least 3 cells, and 3) cells must have at least 250 genes expressed. Additionally, mitochondrial and ribosomal genes were filtered out at this stage. During the second round, the filters focused on the cell subset corresponding to a cell type of interest. With this cell subset, we required a minimum of 10% of the cells to be expressed in either group for the contrast of interest, a minimum of 3 cells per subject, and a minimum of 2 subjects per group.

### Loading of pre-computed DEGs

The pre-computed DEGs can be visualized in cellxgene VIP as shown in Figure 4b and 4d. The user needs to create a sqlite database file which must be in the same folder as the h5ad file and share the same name but with ‘db’ as the file extension, e.g., “ms_nature_2019_Schirmer.db” and “ms_nature_2019_Schirmer.h5ad” are in the same folder to be loaded. If the database file has changed, the server doesn’t need to restart.

There is a script named “DEG2sqlite3.py” under the bin folder, which saves all pre-computed DEG results in csv format to sqlite3 file. It takes 2 parameters where the first parameter is the full path of all csv files, and the second parameter is the name of the database file that will be saved in the same folder as csv files, e.g.,

*python DEG2sqlite3.py ./csv Ms_nature_2019_Schirmer*

The csv files need to contain the 4 columns with headers: gene name, log2FC, Pvalue, FDR. The script is hard coded to pull the first 4 columns in that order. The names of the csv files must be A.vs.B_tag1_tag2.csv, where the contrast is between A and B, and tags can be cell types, e.g., NAWM.vs.NWM_Astrocytes.csv.

### Visualization of spatial transcriptomics data

For pre-processing, we used Scanpy “read_visium” function to read data of each sample from 10x genomics Space Ranger output directory. To note, in order to have the right orientation (Visium capture image border marker on the bottom right should be an empty circle) in the user interface, spatial coordinate Y axis needs to have negative values to fit coordinates design of cellxgene. Therefore, in Python environment such as Jupyter notebook, when plotting the H&E image and coordinates together, they do NOT overlap. On the other hand, when loaded into cellxgene VIP, they would. We also added an additional capability to merge multiple spatial transcriptomics samples into one spatial embedding in a grid format. Our demo example is a 2 x 2 image merged by 4 spatial transcriptomics samples. We apply AnnData.concatenate function to merge 4 multiple AnnData objects into a single AnnData object, then add back individual spatial coordinates stored in AnnData.obsm variable with name starting as “X_spatial_” and original spatial images. After merging, we perform basic QC, apply normalization, log1p transformation, select highly variable genes (HVGs), run PCA dimension reduction, generate UMAP layout and cluster spots by Leiden^60^. At the same time, H&E images were stitched by PIL^61^ Python module and then saved back to Anndata.uns[‘spatial’] as the merged spatial embedding. In summary, all these steps are performed by the below command.

*python3 st_sample_merge.py -i <inputfile of data folders= -o <outputfile> [-d <grid dimension>] [-s <grid cell size=]*

Where, each line in the input file holds one directory in which Space Ranger outputs including aligned images of one Visium slide are stored. Both grid dimension and cell size parameters are optional. Grid dimension is defined in a row by column format (e.g., 2×3 without space) and calculated by default to fit slides in a rectangular grid. The default grid cell size is 700 that is big enough to contain a 600-pixel low resolution image, which is used to minimize the size of the merged image.

Spatial coordinates are visualized by cellxgene embedding layout function, which is normally used for PCA, tSNE or UMAP. To accommodate individual samples in the merged h5ad file, we need to use the same number of spatial spots from the merged samples for each individual sample in the spatial layout, so that any spots that do not belong to an individual sample are assigned (0,0) spatial coordinates, which are displayed as a single dot on the top left corner of the spatial layout with minimum visibility. The user could use a selecting subset cell function from cellxgene to exclude spots not belonging to the sample of interest before plotting or performing analysis. VIP provides a function to retrieve the H&E images and overlay with spatial coordinates. The user can choose either the image or embedding to be on the top layer, adjust transparency, zoom in and out with image and embedding together or separately, realign image and embedding. Further, to facilitate the user to compare and analyze multiple samples together, we allow one to design the desired orientation of samples by arranging, flipping, rotating images of interest into a N x M grid. Once the design is done, the user can download the layout file in JSON format. This JSON file can be supplied to an analyst to generate a new h5ad file with the desired layout of samples by calling the Python script that is provided in the bin directory of the GitHub repository.

*python3 st_h5ad_image_operation.py -j design.json -i merge.h5ad -o final.h5ad*

Once a new h5ad file is created, an analyst can spin up another cellxgene VIP instance of the finalized data set for the user to explore. This workflow encourages close collaboration between experimentalists and analysts and prevents unintended large h5ad files from proliferating if an end user is given an option to generate such files on the fly through the user interface.

### Command line interface (CLI)

On client side, we embed the Ace editor (https://ace.c9.io) in VIP for advanced users to write analysis code to operate on either the whole dataset or a smaller sliced data set based on selections of cells and genes. The editor offers syntax coloring, automated indent, highlighting matching parentheses and other advanced features to match performance of native editors such as Vim. For Python programmers, Jupyter notebook syntax is fully supported, including built-in magic commands, e.g., to set the output image format to SVG.

*%config InlineBackend.figure_formats = [‘svg’]*

The browser then passes the code written in the CLI editor to the server side, where the following one-line command firstly converts the Python script submitted from the client to Jupyter notebook document by “jupytext” (https://jupytext.readthedocs.io) and secondly runs “jupyter nbconvert” (https://nbconvert.readthedocs.io) to execute the notebook and generate output in HTML format that will be returned to the client side.

*jupytext --to notebook --output - <input script> | jupyter nbconvert --ExecutePreprocessor.timeout=1800--to html --execute --stdin –stdout*

To communicate with big data hosted on server side in CLI, two preserved variables are made visible to both the client and the server side. One is “adata” that is a Python AnnData object holding gene expression matrix of the whole dataset or selected cells and metadata while the other is “strPath” that is the file name including path but leaving out the file extension. Thus, concatenation of strPath and “.h5ad” is the name of the file storing “adata” that is derived from the whole data based on user selections and stored on the server temporally for use in a session executing the script submitted from the interface.

To be flexible, there are two ways of programming in R. One is using R magic enabled by rpy2 module that is already installed on server side. As shown in CLI Vignette 2, the following code snippet shows how to use apache Arrow technique to share “adata” object in feather file format (https://bit.ly/3dOi0Lu) between Python and R code blocks.

*from pyarrow import feather*

*import pandas as pd*

*X = adata.to_df()*

*meta = pd.concat([adata.obs,adata.obsm.to_df()],axis=1)*

*feather.write_feather(X,strPath+‘.feather’)*

*feather.write_feather(meta,strPath+“.obs.feather”)*

*%%R*

*D <- t(arrow::read_feather(paste(strPath,“.feather”,sep=“”)))*

*meta <- as.data.frame(arrow::read_feather(paste(strPath,“.obs.feather”,sep=“”)))*

*colnames(D) <- meta[,“name_0”]*

*X <;- CreateSeuratObject(counts=D)*

*X@meta.data <- cbind(X@meta.data,meta)*

As illustrated in CLI Vignette 3, R markdown is another way to program in R. The reticulate package is used to run Python code in R markdown to prepare the data file the same way as above. On the server side, the following command is executed to generate the returned html report.

*Rscript -e ‘rmarkdown::render(“<input R markdown>”, output_file=“output.html”)’*

All Python modules and R packages used in CLI environment need to be installed beforehand by following the installation instruction in the GitHub.

### Visualization of multiome data

Cellxgene VIP supports the visualization of multiome data to illustrate the relationship of DNA accessibility and expression of nearby genes at a locus. To exemplify this functionality, we used a PBMC multiome dataset from 10x Genomics^46^. For pre-processing, we utilized Signac^20^ and Seurat to make a joint RNA and ATAC analysis with the hg38 reference genome. DNA accessibility data was used to compute per-cell quality control metrics and filter out the low-quality cells. After quality control, gene expression data was processed by SCTranfrom^62^ and PCA while DNA accessibility data was processed by peak-calling, latent semantic indexing, and single value decomposition. After normalization and dimension reduction, we used a curated Azimuth human PBMC reference^17^ to annotate cell types through Seurat reference mapping. To visualize the joint embedding plot, the weighted nearest neighbor method and UMAP were applied.

Moreover, with scATAC-seq data, VIP allows users to visualize track plot in a mini genome browser. In Signac, we can calculate the correlation between gene expression and accessibility at the nearby peaks by using LinkPeaks to compute the GC content, and linkage between peaks and genes. The frequency plot over genomic region, coordinates, peaks distribution and links can be visualized by the CoveragePlot in Signac. However, fetching peaks and links annotation from the Seurat object which contains the whole genome information and displaying them in real-time on VIP interface are challenging. As a viable alternative, we used the annotated cell types to split scATAC bam files by Sinto^63^ and made the BIGWIG files separately by bamCoverage in deepTools^64^. Links, peaks, and genomic range information from Signac session were saved as ‘links.rds’, ‘annotation.rds’ and ‘peaks.rds’. All files with ‘rds’ and ‘bw’ extension are required to be in a folder with the same names as the corresponding h5ad file holding scRNA-seq data. In addition, it also requires a file named “bw.cluster” in the folder. The first row in “bw.cluster” is the metadata category which BIGWIG files were split on; for example, we split on ‘CellType’ category with PBMC multiome dataset. The following two columns should indicate the filenames of bigwigs, and corresponding categorical names in h5ad. Upon the availability of these files, “Multiome” menu in VIP will be activated when such dataset is loaded in cellxgene VIP. The detailed reproductible process of the demo data is available at “Data Availability” section.

### Tutorial

A comprehensive user guide to illustrate detailed functionality of each module with step-by-step instruction is available as supplementary information and also accessible conveniently as an online HTML Bookdown^65^ document at https://interactivereport.github.io/cellxgene_VIP/tutorial/docs. Inside the application, a question mark behind the title of each function module provides a direct way to reach the corresponding section of the HTML document for help.

### Demos

sWe demonstrate the functionalities and capabilities of VIP on visualizing scRNA-seq, spatial transcriptomics, and multiome datasets by the following links, https://cellxgenevip-ms.bxgenomics.com, https://cellxgenevip-spatial.bxgenomics.com, and https://cellxgenevip-multiome.bxgenomics.comrespectively.

## Code availability

The source code and detailed installation guide are provided at https://github.com/interactivereport/cellxgene_VIP. Cellxgene VIP is released under the MIT License.

## Data availability

snRNA-seq. The demo MS snRNA-seq data in h5ad format can be reproduced by following the Jupyter notebook https://bit.ly/2CeUHtO or downloaded from https://bit.ly/3JjQqSp^66^.

Spatial transcriptomics. 4 Visium Spatial Transcriptomics datasets were downloaded from the 10X Genomics data site and were reformatted as spaceranger outputs for the convenience of reproducible downstream analysis. Data^67^ can be downloaded from https://bit.ly/3uut88d. The demo h5ad file can be reproduced by following the notebook https://bit.ly/3O1fyRh or by this command, “python3 st_sample_merge.py -i inputfiles.txt -o merged_demo.h5ad -d 2×2 -s 600”.

Multiome. The publicly available 10x Genomic Multiome dataset can be downloaded from https://bit.ly/3urczd4. The demo multiome h5ad and bigwig files can be reproduced by following https://bit.ly/3LXnUaD.

## Acknowledgements

The authors are grateful and indebted to the cellxgene team from Chan Zuckerberg Initiative for the development of the base framework. We would also like to thank Dr. L. Schirmer and Prof. D.H. Rowitch for allowing us to use scRNA-seq data on Multiple Sclerosis for the demonstration of cellxgene VIP.

## Author contributions

K.L, Z.O. and B.Z. designed and implemented the software. All authors contributed to testing of the tool. K.L., Y.C., J.G., D.L., M.M. and B.Z. wrote the manuscript with input from all authors. All authors read and approved the manuscript.

