## Supplementary file for "Cellxgene VIP unleashes full power of interactive visualization and integrative analysis of scRNA-seq, spatial transcriptomics, and multiome data"

### 1 Getting started with cellxgene VIP

This is a cellxgene VIP supplementary information including a tutorial book written in **Bookdown**.

#### 1.1 Why use cellxgene VIP?

To meet the growing demands from scientists to effectively extract deep insights from single cell RNA sequencing, spatial transcriptomics, and emerging multiome datasets, we developed cellxgene VIP (Visualization In Plugin), a frontend interactive visualization plugin of cellxgene framework, which greatly expanded capabilities of the base tool in the following aspects. First, it generates a comprehensive set of over eighteen commonly used quality control and analytical plots in high resolution with highly customizable settings in real time. Second, it provides more advanced analytical functions to gain insights on cellular compositions and deep biology, such as marker gene identification, differential gene expression analysis, and gene set enrichment analysis. Third, it empowers advanced users to perform analysis in a Jupyter Notebook like environment, dubbed Command Line Interface (CLI) by programming in Python and/or R directly without limiting themselves to functional modules available via graphical user interface (GUI). Finally, it pioneers methods to visualize multi-modal data, such as spatial transcriptomics embedding aligned with histological image on one slice or multiple slices in a grid format, and the latest 10x Genomic Multiome dataset where both DNA accessibility and gene expression in the same cells are measured, under the same framework in an integrative way to fully leverage the functionalities mentioned above. Taken together, the open-source tool makes large scale single cell data visualization and analysis more accessible to biologists in a user-friendly manner and fosters computational reproducibility by simplifying data and code reuse through the CLI. Going forward, it has the potential to become a crowdsourcing ecosystem for the scientific community to contribute even more modules to the Swiss Army knife of single cell data exploration tools.

#### 1.2 Getting Set up

##### 1.2.1 Execute anaconda

```
bash ~/Downloads/Anaconda3-2020.02-Linux-x86_64.sh
```

If anaconda is not installed on server, you can install it following anaconda documentation (https://docs.anaconda.com/anaconda/install/linux/) ### Create and enable conda environment

```
# clone repo from cellxgene VIP github
git clone https://github.com/interactivereport/cellxgene_VIP.git
cd cellxgene_VIP

# conda environment
source <path to Anaconda3>/etc/profile.d/conda.sh (Default:
↪  /opt/anaconda3/etc/profile.d/conda.sh)
conda config --set channel_priority flexible
conda env create -n <env name, such as: VIP> -f VIP.yml (system-wide R) or
↪  VIP_conda_R.yml (local R under conda, no root privilege needed)
```

Activate conda environment

```
conda activate <env name, such as: VIP>
```

or

```
source activate <env name>
```

##### 1.2.2 Cellxgene installation

Install cellxgene by running config.sh in "cellxgene_VIP" directory

```
./config.sh
```

##### 1.2.3 R dependencies

Install all required R packages on linux:

```
export LIBARROW_MINIMAL=false
#  ensure that the right instance of R is used. e.g. system-wide: /bin/R or /usr/bin/R
↪  ; local R under conda: ~/.conda/envs/VIP_conda_R/bin/R
which R

R -q -e 'if(!require(devtools)) install.packages("devtools",repos =
↪  "http://cran.us.r-project.org")'
R -q -e 'if(!require(Cairo)) devtools::install_version("Cairo",version="1.5-12",repos
↪  = "http://cran.us.r-project.org")'
R -q -e 'if(!require(foreign))
↪  devtools::install_version("foreign",version="0.8-76",repos =
↪  "http://cran.us.r-project.org")'
R -q -e 'if(!require(ggpubr)) devtools::install_version("ggpubr",version="0.3.0",repos
↪  = "http://cran.us.r-project.org")'
R -q -e 'if(!require(ggrastr))
↪  devtools::install_version("ggrastr",version="0.1.9",repos =
↪  "http://cran.us.r-project.org")'
R -q -e 'if(!require(arrow)) devtools::install_version("arrow",version="2.0.0",repos =
↪  "http://cran.us.r-project.org")'
R -q -e 'if(!require(Seurat)) devtools::install_version("Seurat",version="3.2.3",repos
↪  = "http://cran.us.r-project.org")'
R -q -e 'if(!require(rmarkdown))
↪  devtools::install_version("rmarkdown",version="2.5",repos =
↪  "http://cran.us.r-project.org")'
R -q -e 'if(!require(tidyverse))
↪  devtools::install_version("tidyverse",version="1.3.0",repos =
↪  "http://cran.us.r-project.org")'
R -q -e 'if(!require(viridis))
↪  devtools::install_version("viridis",version="0.5.1",repos =
↪  "http://cran.us.r-project.org")'
R -q -e 'if(!require(BiocManager))
↪  devtools::install_version("BiocManager",version="1.30.10",repos =
↪  "http://cran.us.r-project.org")'
R -q -e 'if(!require(fgsea)) BiocManager::install("fgsea")'

# These should be already installed as dependencies of above packages
R -q -e 'if(!require(dbplyr)) devtools::install_version("dbplyr",version="1.0.2",repos
↪  = "http://cran.us.r-project.org")'
R -q -e 'if(!require(RColorBrewer))
↪  devtools::install_version("RColorBrewer",version="1.1-2",repos =
↪  "http://cran.us.r-project.org")'
R -q -e 'if(!require(glue)) devtools::install_version("glue",version="1.4.2",repos =
↪  "http://cran.us.r-project.org")'
R -q -e 'if(!require(gridExtra))
↪  devtools::install_version("gridExtra",version="2.3",repos =
↪  "http://cran.us.r-project.org")'
```

```
R -q -e 'if(!require(ggrepel))
↪  devtools::install_version("ggrepel",version="0.8.2",repos =
↪  "http://cran.us.r-project.org")'
R -q -e 'if(!require(MASS)) devtools::install_version("MASS",version="7.3-51.6",repos
↪  = "http://cran.us.r-project.org")'
R -q -e 'if(!require(data.table))
↪  devtools::install_version("data.table",version="1.13.0",repos =
↪  "http://cran.us.r-project.org")'
```

###### 1.2.4 Run cellxgene by h5ad file

You can aslo run cellxgene by specifying a h5ad file, which stores scRNA-seq data along with a host and a port. Use 'ps' to find used ports to spare. Please see https://chanzuckerberg.github.io/cellxgene/posts/launch for details

```
ps -ef | grep cellxgene
Rscript -e 'reticulate::py_config()'
# Run the following command if the output of the above command doesn't point to the
↪  Python in your env.
export RETICULATE_PYTHON=`which python`
cellxgene launch --host <xxx> --port <xxx> --disable-annotations --verbose <h5ad file>
```

###### 1.2.5 Cellxgene on web browser

chrome is preferred, version 87.0.4280.88 or 87.0.4280.141 is used. Users can access **http(s)://host:port**. Following screenshot is what you should be able to see in console of chrome developer tools.

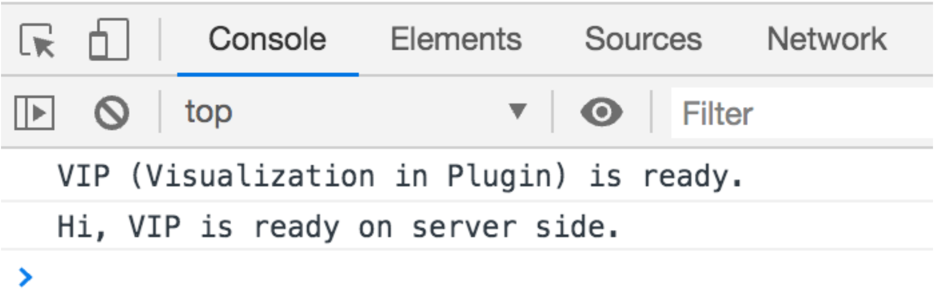

##### 1.3 Authors

- Keijie Li, Associate Director at Biogen in the research department. Main author and content wrangler.
- Zhengyu Ouyang, Associate Director of Bioingormatics at BioinfoRx. Content wrangler.

The development and delivery of this material has also contributed by:

- Yirui Chen, Biogen Inc.
- Dongdong Lin, Biogen Inc.
- Michael Minguenau, Biogen Inc.
- Jake Gagnon, Biogen Inc.
- Will Chen, Biogen Inc.
- David Sexton, Biogen Inc.

#### 2 How to use Cellxgene VIP

#### 2.1 Graphical user interface of cellxgene and VIP

The main window of cellxgene is divided into three regions, the left panel mainly displays categorial annotations, brief description of the data set and initial graphics setting, specifically embedding and coloring of cells. On the right panel, it hosts continuous variables, such as qc metrics shown in histogram with x, y corresponding to values of a measurement and numbers of cells, respectively. More importantly, cells shown as individual dots are presented in the center panel based on a selected embedding and colored by either categorial annotations or continuous variables, which is indicated by pressed rain drop icon, e.g., cell_type in Supplementary Fig. 1a.

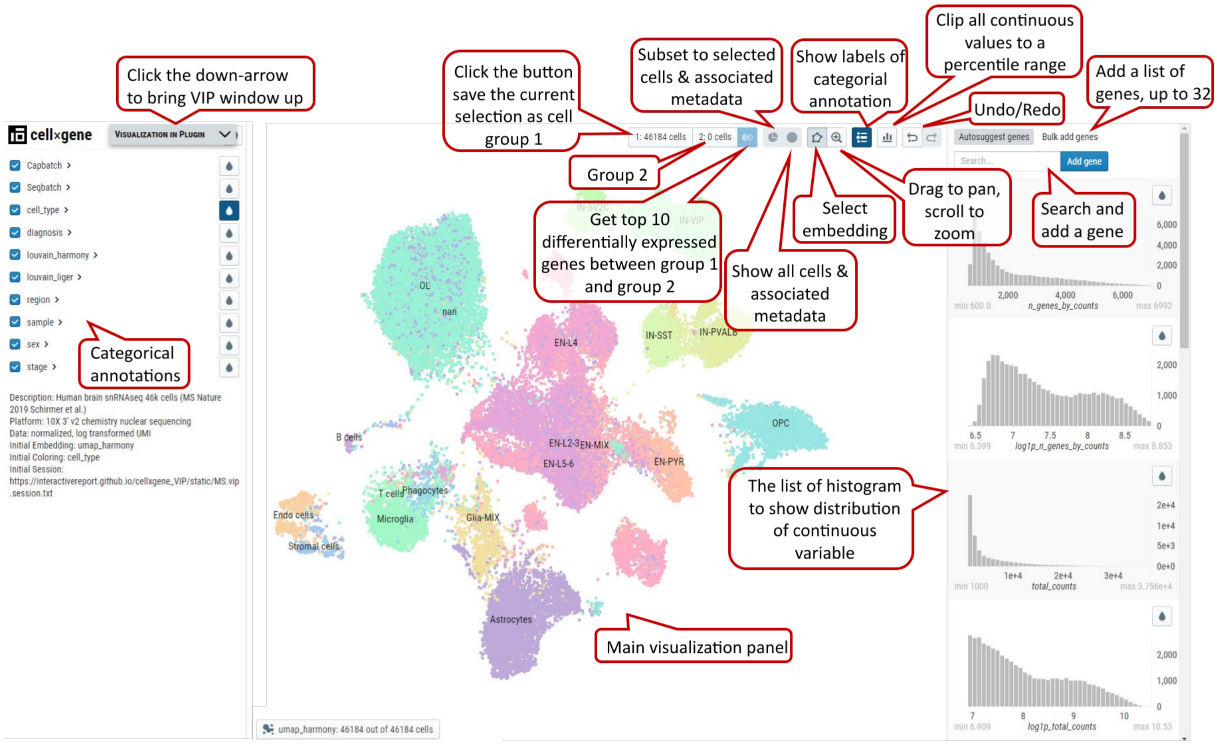

Supplementary Fig. 1a. Cellxgene main window, functional icons and minimized VIP bar next to cellxgene logo.

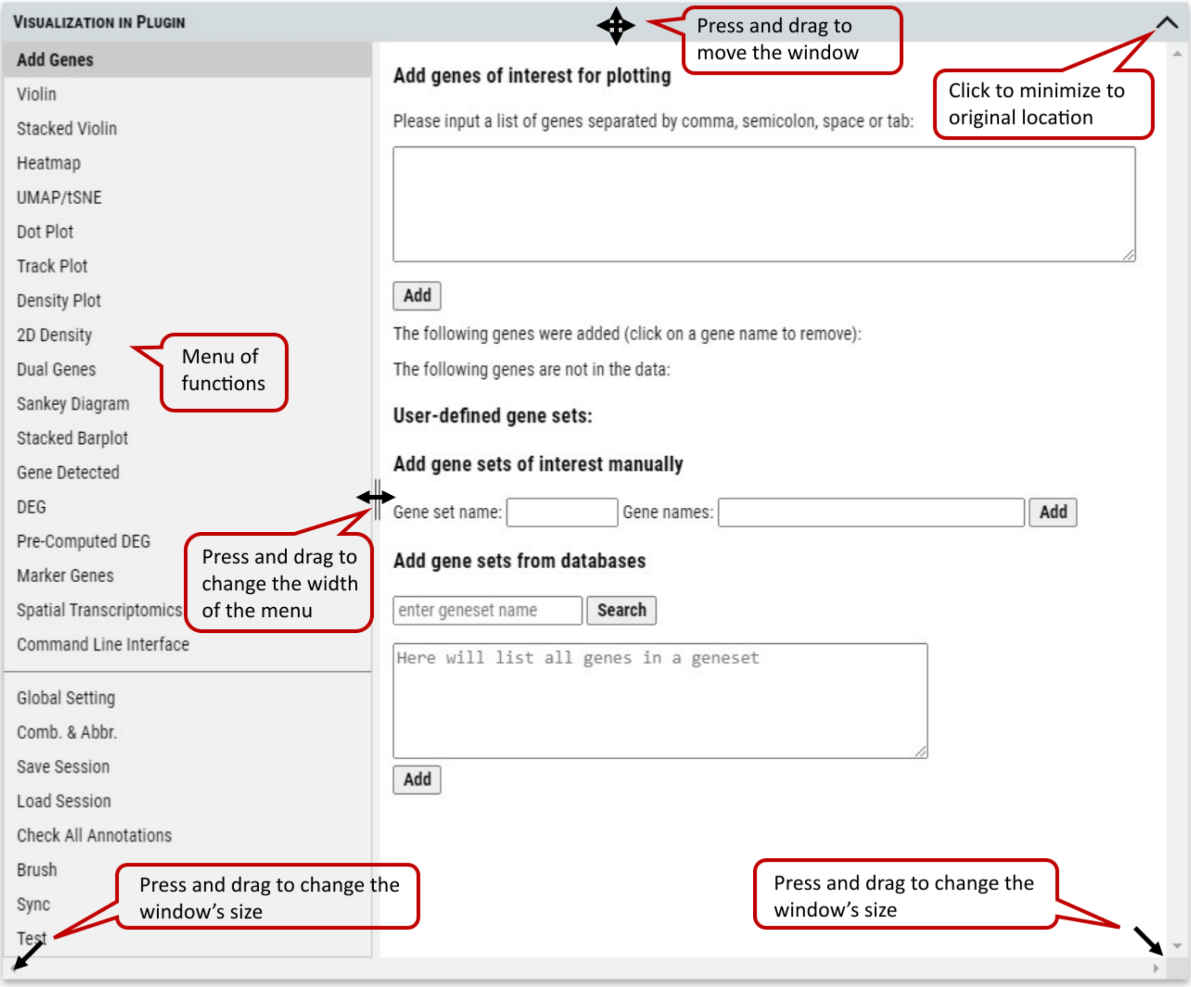

Supplementary Fig. 1b. VIP (Visualization in Plugin) window and controls of user interface. The cursor will change to corresponding icon when mouse hovers over control anchors inside the window. In the case of missing title bar after operation, changing the size of outside browser window (not VIP window) will always bring the VIP window back to the original location near the cellxgene logo.

#### 2.2 Cell selection by categorial annotations

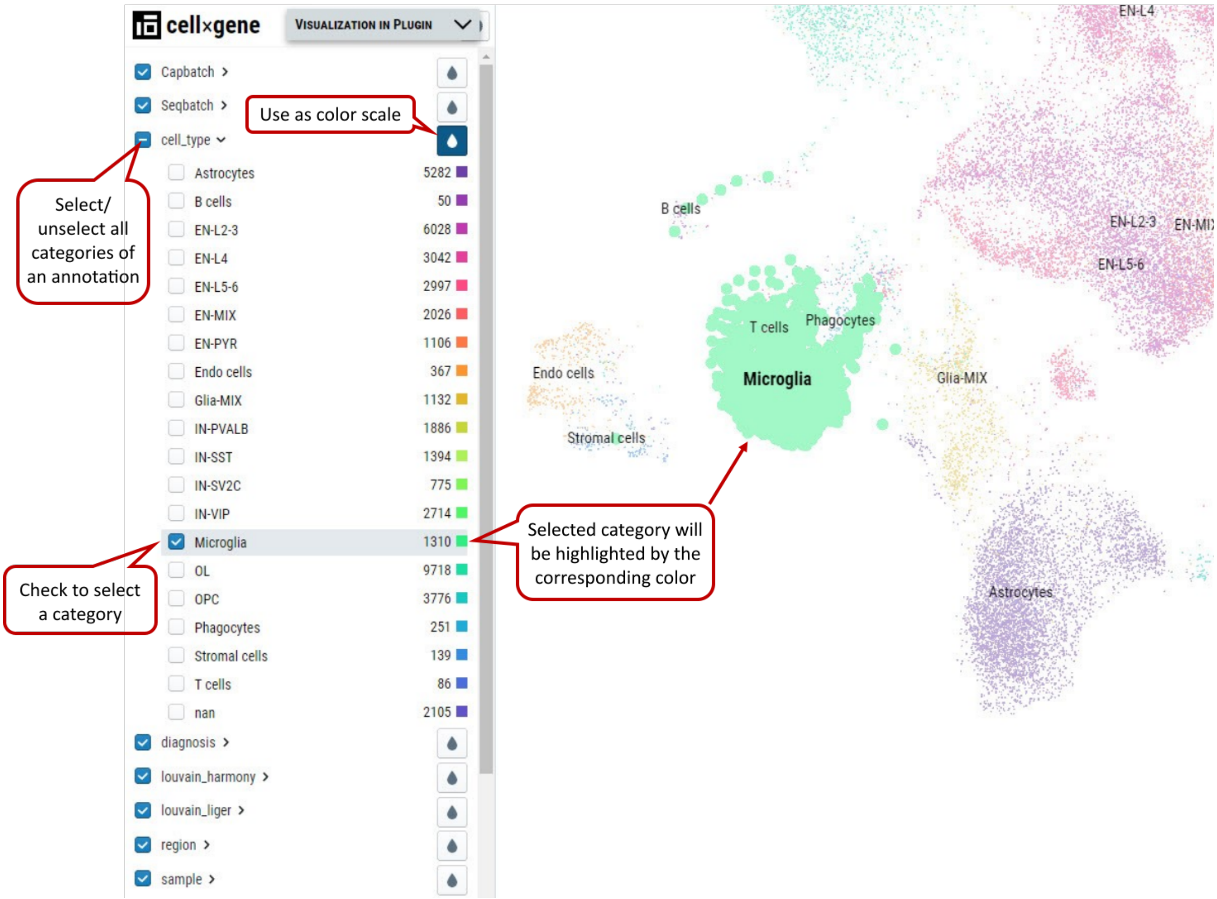

Supplementary Fig. 2. Cell selection by categorial annotation. Selected B cells are shown in bold dots and highlighted in purple color when hovering the mouse over the cluster.

It is an overlap operation when categories from multiple annotations are checked to make the final selection. E.g., if male from sex is also checked besides B cells, it means cells from B cells cluster of male samples are selected. Note: Click “1:” or “2:” button to save cell selection into group 1 or 2

#### 2.3 Cell selection by brushing on distribution of continues variables

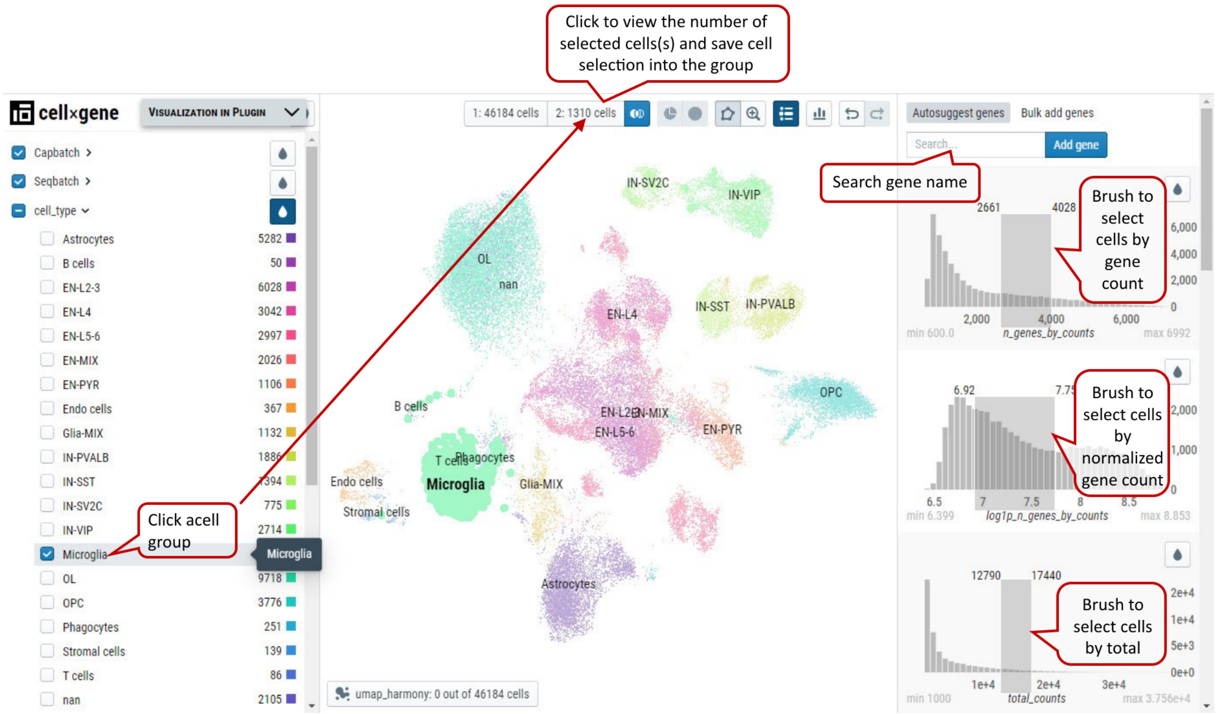

Supplementary Fig. 3. Cell selection by brushing the ranges of continuous variables. Low- and high-end values are shown at the top corners of the brushing boxes in dark gray.

Note: Histograms of expression values of genes can by brushed as well to get cells expressing certain genes in the range.

#### 2.4 Free hand Lasso selection on dots representing cells

From the cell visualization panel, user can freely select a cluster of cells of interest by using 'Lasso' selection tool. The selected cluster of cells can also be added as a group for downstream analysis in cellxgene VIP.

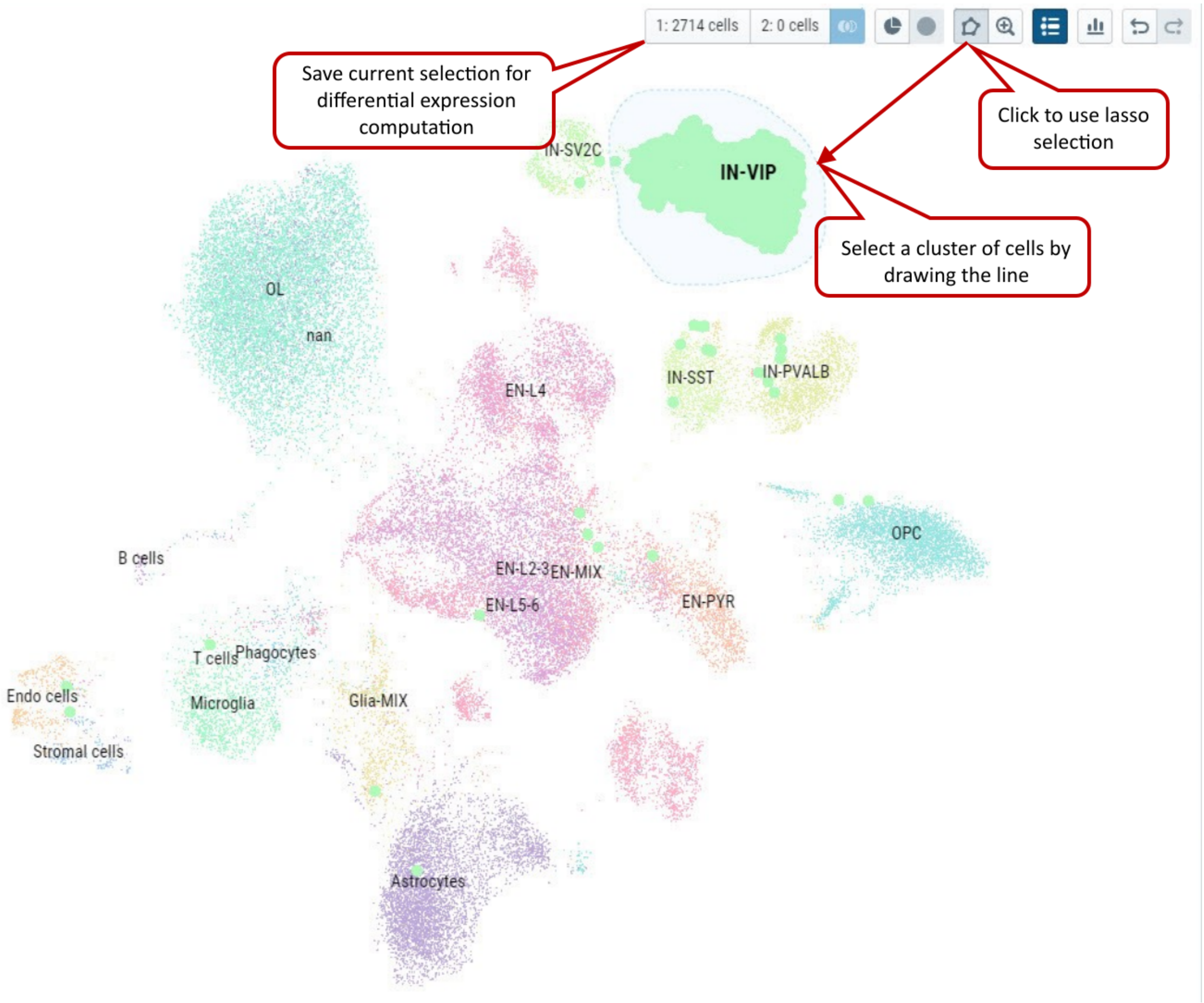

Supplementary Fig. 4. Select cells by using free hand Lasso selection tool and add these cells as a group for further analysis in cellxgene VIP.

Note: Please try to draw as close as possible to the starting point in the end to make an enclosed shape to ensure successfully lasso selection.

#### 2.5 VIP – Global Setting

User can set parameters for figure plotting that control plotting functions except CLI. 'split_show' branch of Scanpy offers better representation of Stacked Violin and Dot Plot comparing to master branch.

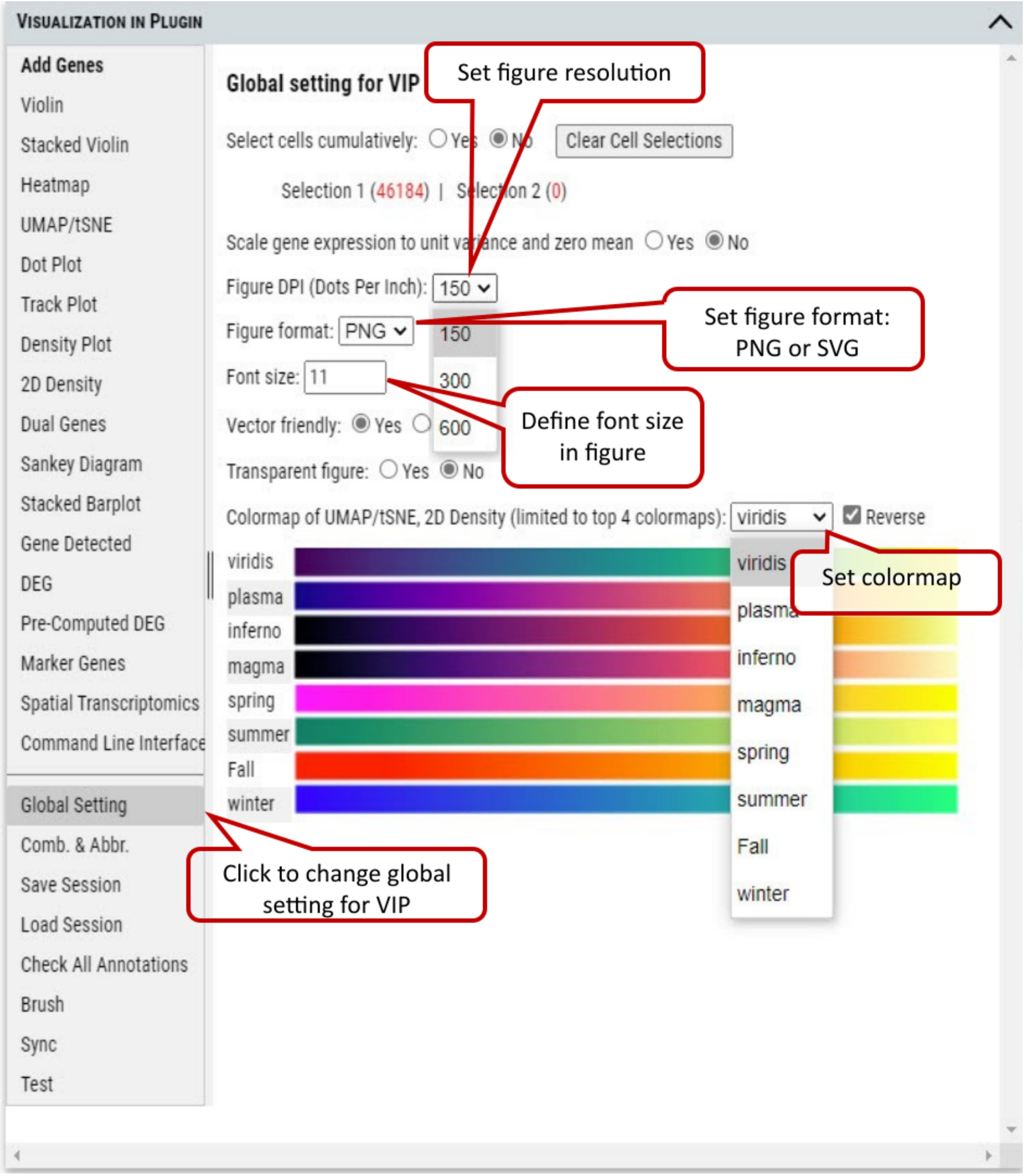

Supplementary Fig. 5. Setting parameters for figure plotting.

Scaled data have zero mean and unit variance per gene. This was performed by calculating z-scores of the expression data using Scanpy's scale function. (Scanpy pp.scale function: Scale data to unit variance and zero mean.)

We provide flexibility to allow 1) scale to unit variance or not; 2) Zero centered or not; 3) Capped at max value after scaling.

We recommend using scaled data for plotting/visualization while using non-scaled data for differential gene expression analysis.

Note: Dot plot is one exception in visualization category which uses non-scaled data for meaningful interpretation.

#### 2.6 VIP – Add Genes / Gene Sets

Cellxgene VIP allows user to add any genes or gene sets for extensive exploration and visualization. User can either type a list of gene in the textbox or create sets of genes to be grouped together in plots. Then the genes will be automatically listed for plotting in other functional modules after checking availability in the dataset.

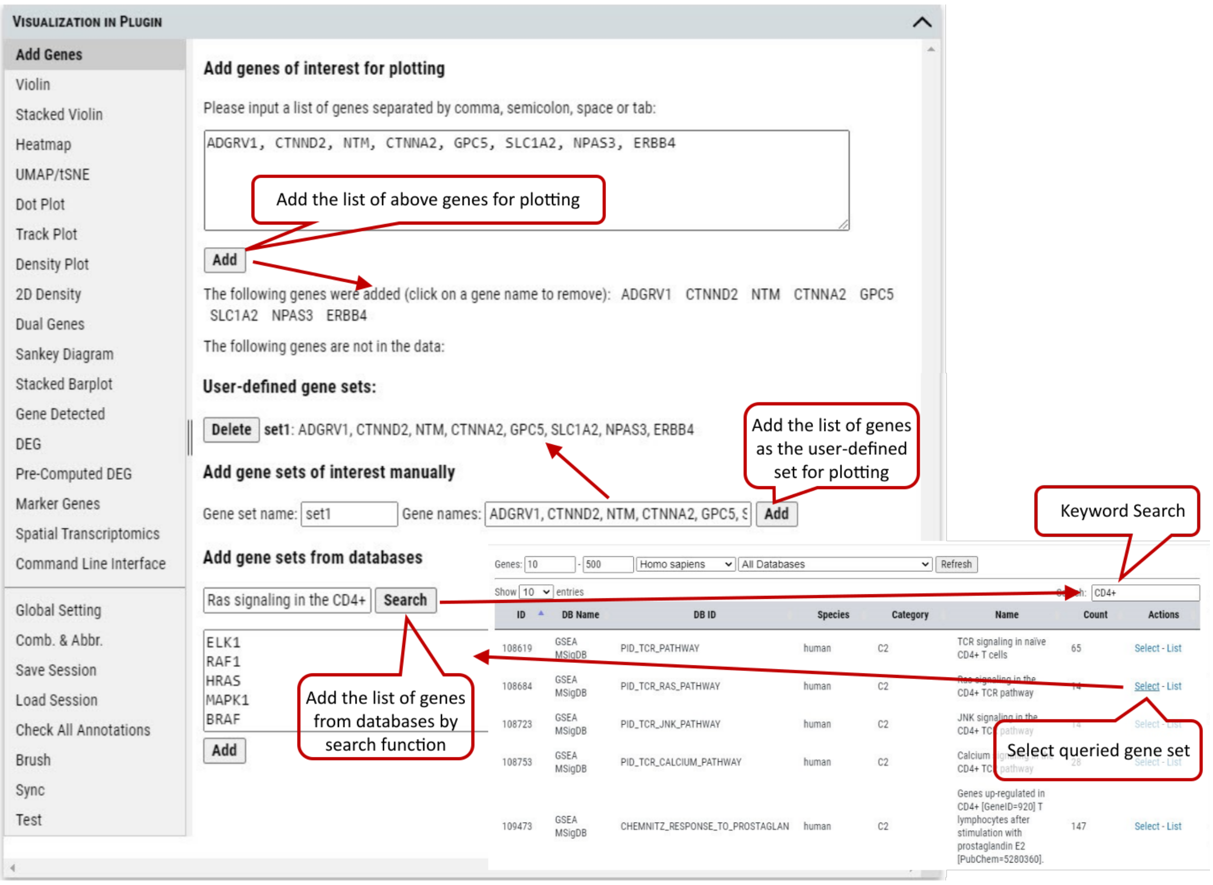

Supplementary Fig. 6. Add gene or gene sets for plotting.

Note: The cursor will turn to cross icon while hovering over a gene name, then click to delete the gene.

#### 2.7 VIP – Violin Plot

To plot expression of gene among categories of an annotation, e.g., cell type, sex, or batch etc. Step 1. User needs to select the group(s) of cells for plotting. These groups can be created by using selection tools illustrated in tutorial section 2, 3 and/or 4. Initially, all of cells are gathered in 'Group 1' by default. Step 2. Select a gene from the gene list which could be added as shown in section 6. An expression level cutoff can be set to further filter out cells with low level expression of such gene. Step 3. Select the annotation to group cells for plotting. Step 4. Execute plotting, get plotting data (i.e., gene expression), manipulate image (e.g., zoom in/out) or save the image.

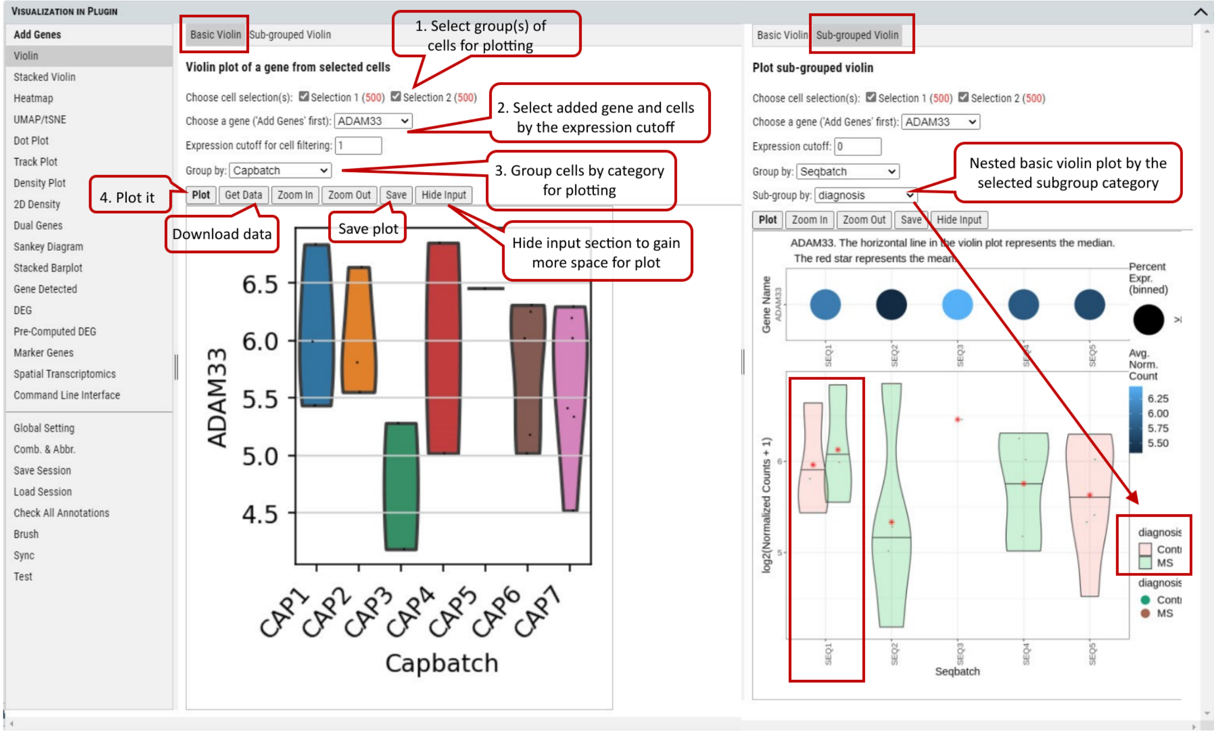

Supplementary Fig. 7. Violin plot of gene expression values of a gene grouped by cell type.

Note: Figure resolution and format can be set in "Global Setting" tab as shown in tutorial section 5.

#### 2.8 VIP – Stacked Violin

Beyond plotting expression values of a gene, stacked violin allows plotting of multiple genes together.

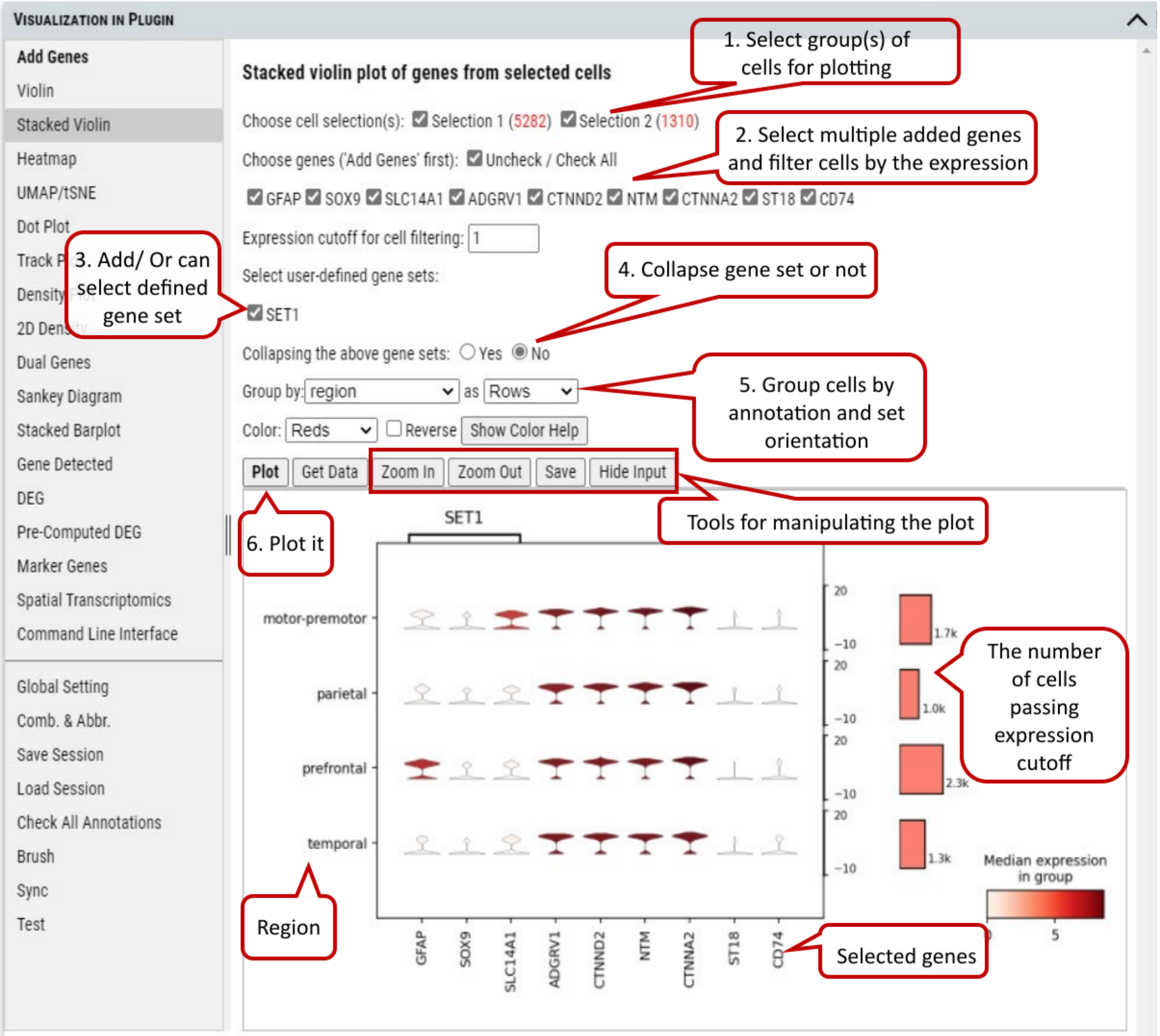

Supplementary Fig. 8. Stacked Violin plot of multiple genes and/or gene set.

Note: If collapsing of gene sets is set to 'Yes', average gene expression of genes in a set is used for plotting.

#### 2.9 VIP – Heatmap

To show or compare the expression level (i.e., expression value or expression Z-score) of multiple genes among the selected group of cells.

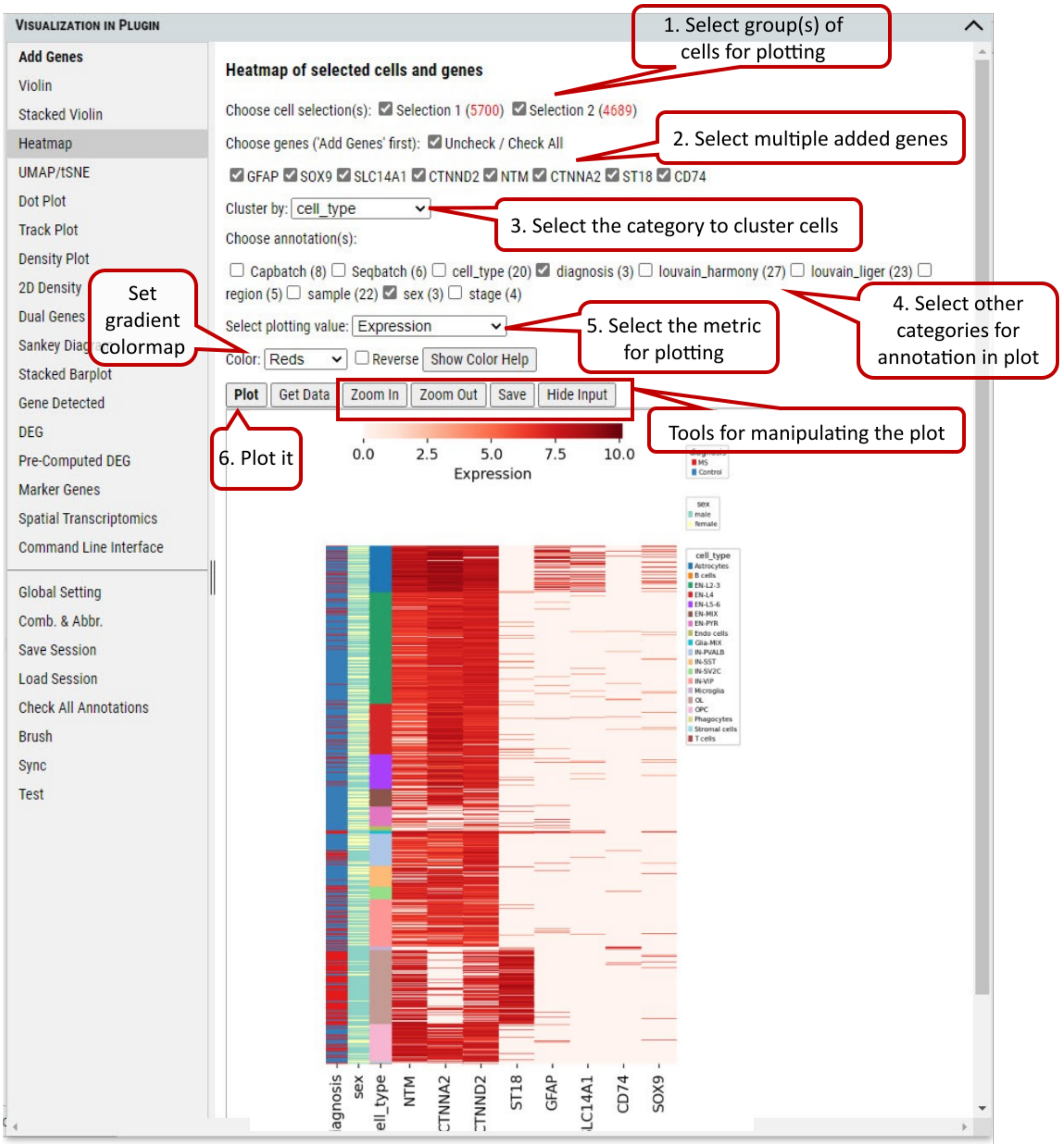

Supplementary Fig. 9. Heatmap of gene expression in cells grouped by annotations.

#### 2.10 VIP – Embedding

To plot the embedding (UMAP, tSNE, or spatial) of cells in the selected group(s). One of pre-computed and loaded embeddings can be selected.

The user can color cells in the embedding plots by multiple annotations (e.g., cell_type, diagnosis).

Besides coloring cells by annotations, user can color cells based on gene expression level of selected genes in the embedding plots.

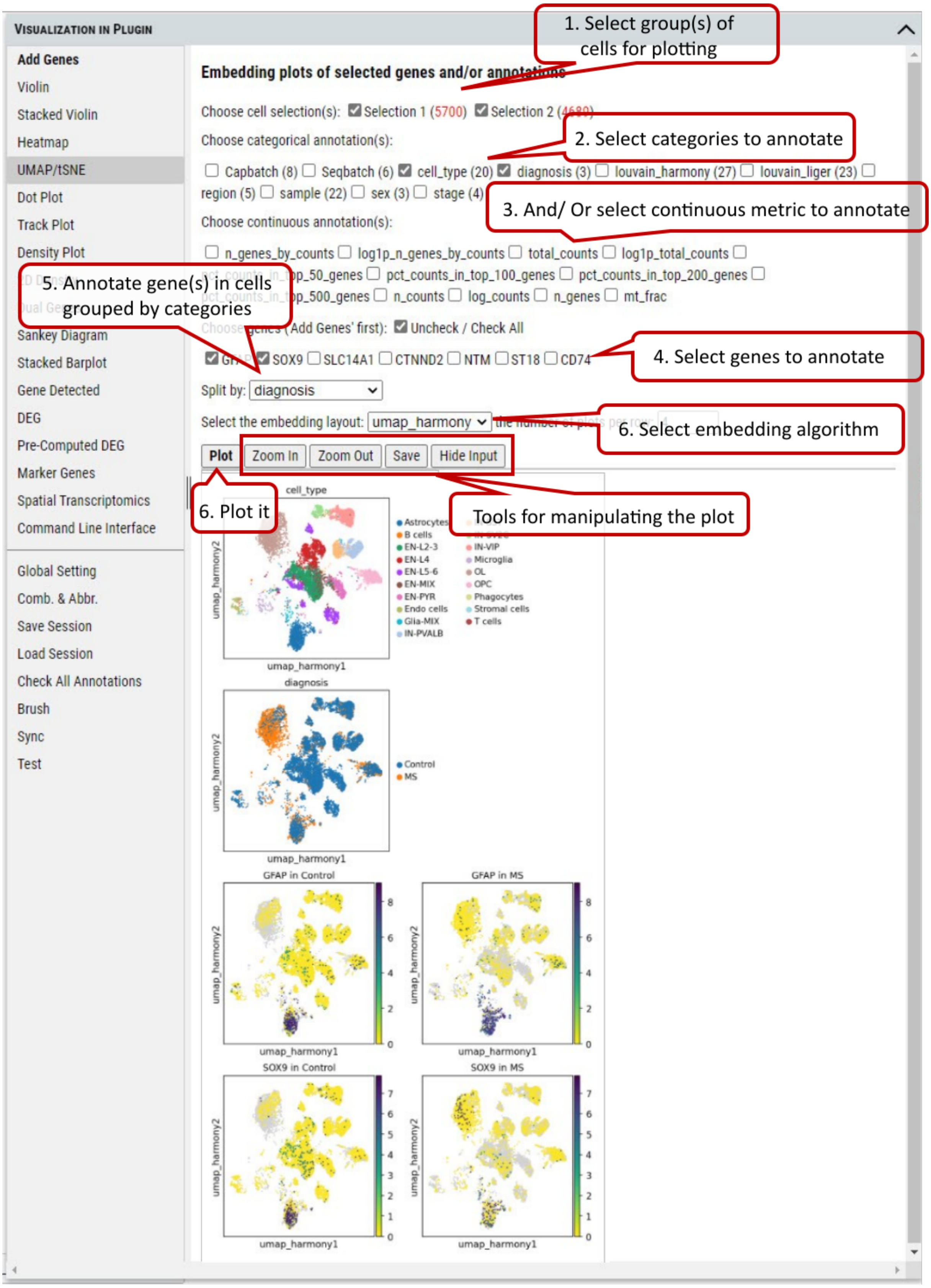

Supplementary Fig. 10. Embedding plotting of expression level of genes or gene set in the cells split by categories of an annotation.

#### 2.11 VIP – Dot Plot

To show the fraction of cells (annotated by dot size) expressing a gene in each group and the averaged expression level of the gene (annotated by color intensity) in the group.

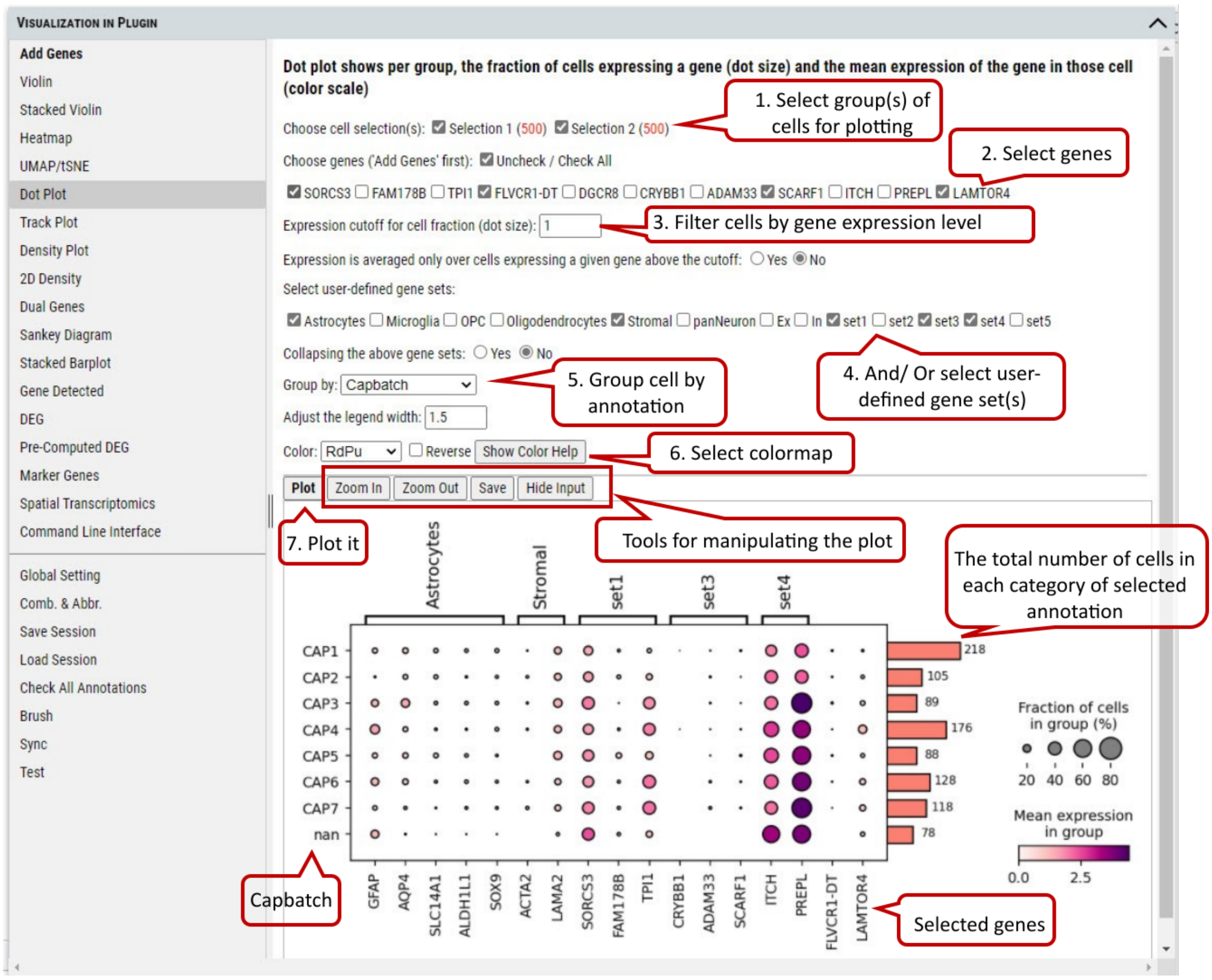

Supplementary Fig. 11. Dot plotting of the fraction of cells expressing genes above a cutoff in each categorie of the selected annotation.

Note: The number of cells represented by side bar chart are always numbers of cells distributed in each category of certain annotation without filtering. It will give an accurate estimate of number of cells in each bubble in the plot. The use of the plot is only meaningful when the counts matrix contains zeros representing no gene counts. Its visualization does not work for scaled or corrected matrices in which zero counts had been replaced by other values, see https://scanpy-tutorials.readthedocs.io/en/multiomics/visualizing-marker-genes.html#Dot-plots.

#### 2.12 VIP – Track Plot

To show the expression of gene(s) of individual cells as vertical lines grouped by the selected annotation on x-axis. Instead of a color scale, the gene expression is represented by height.

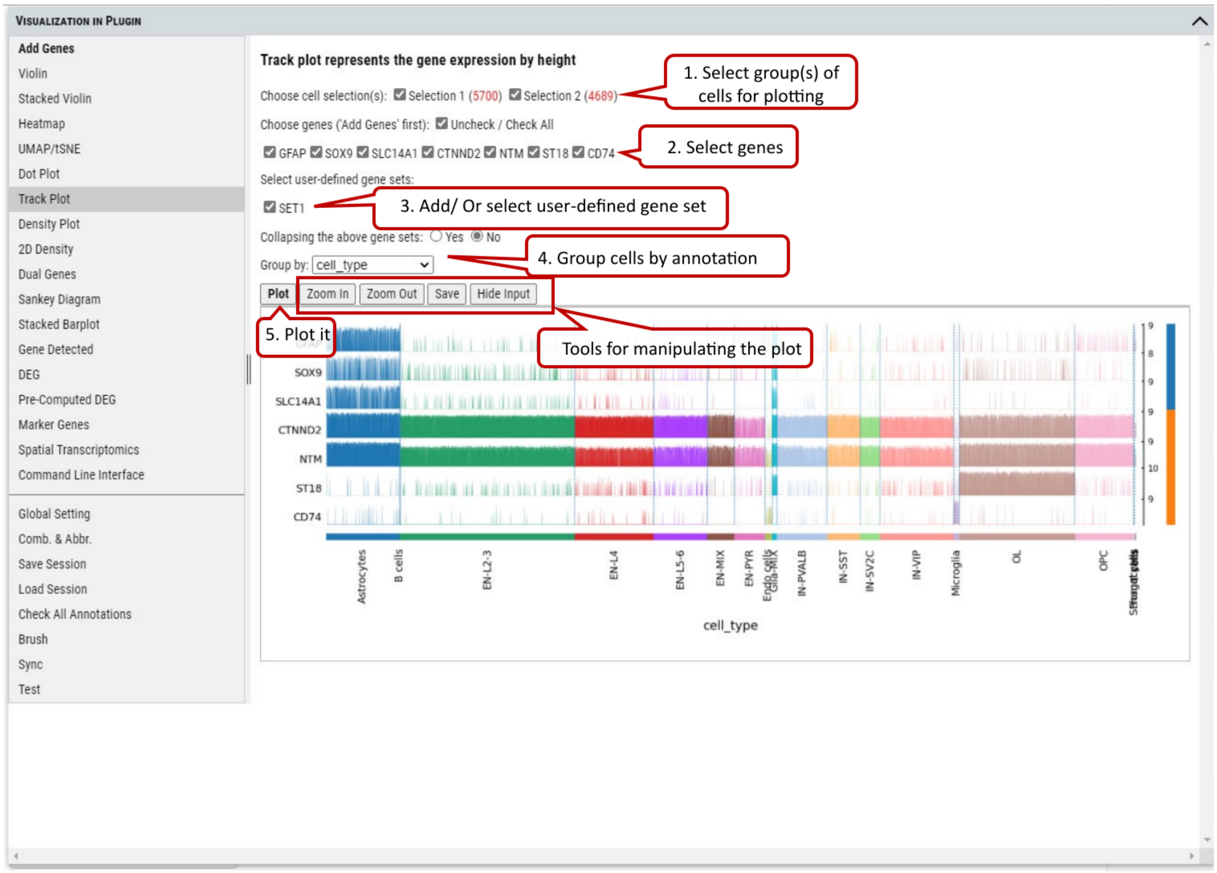

Supplementary Fig. 12. Track plotting of expression of genes or gene set in each category of the selected annotation. Gene expression levels are represented by the heights of vertical lines.

#### 2.13 VIP – Density Plot

To show the density of gene(s) expression in the cells annotated by category in the selected group(s) of cells. A density plot is a representation of the distribution of a numeric variable. It uses a kernel density estimate to show the probability density function of the variable (see more). It is a smoothed version of the histogram and is used in the same concept.

The bandwidth defines how close to a value point the distance between two points must be to influence the estimation of the density at the point. A small bandwidth only considers the closest values, so the estimation is close to the data. A large bandwidth considers more points and gives a smoother estimation.

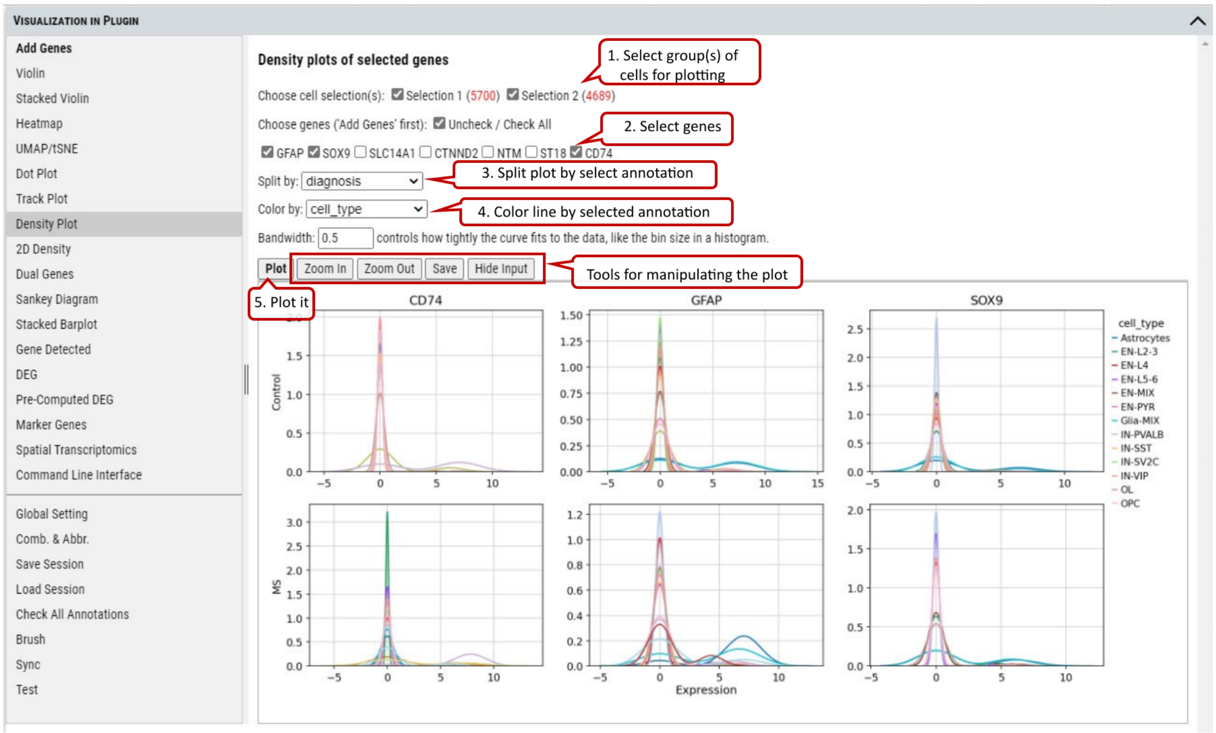

Supplementary Fig. 13. Density plotting of the expression of genes in each group split by one annotation while colored by another.

#### 2.14 VIP – Density Scatter Plot

Besides plotting of expression density of single gene, density scatter plot allows to explore the joint expression density of two genes in the cells expressing both genes above a cutoff.

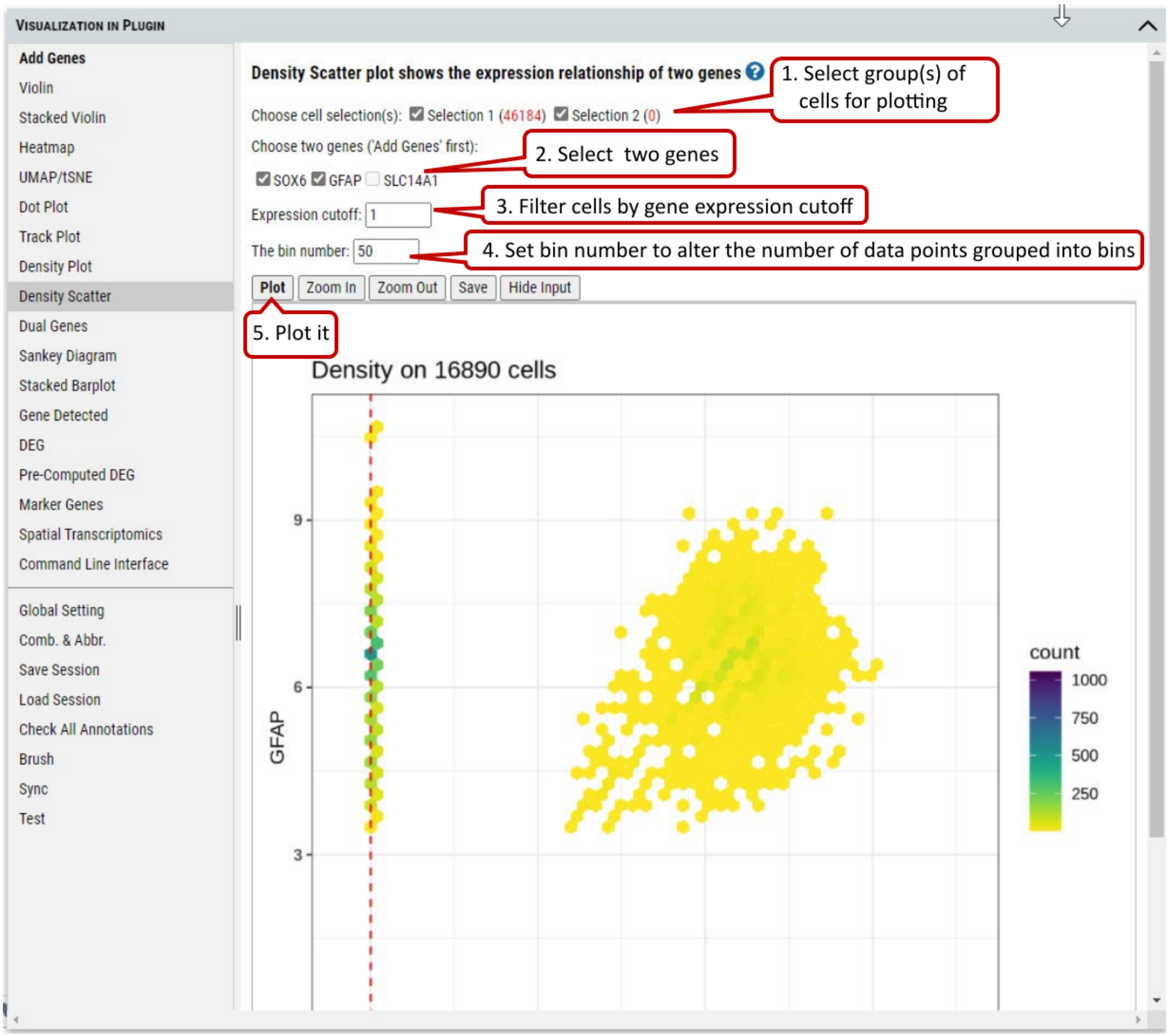

Supplementary Fig. 14. Density scatter plotting of expression of two genes in the selected cells.

#### 2.15 VIP – Dual Genes

To view the relationship of expression levels of two genes in selected cells. It is based on the embedding plot of cells while coloring cells according to the expression levels of gene(s) in each cell.

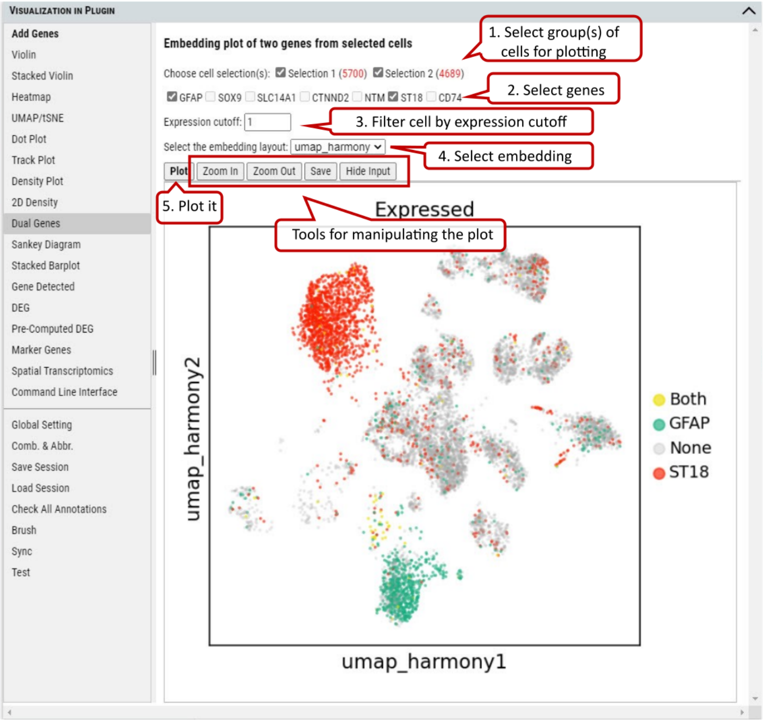

Supplementary Fig. 15. Embedding plotting of the expression of two genes in the selected cell group(s).

#### 2.16 VIP – Sankey Diagram

Sankey diagram shows the flow of gene expression and annotations linked by cells. Gene expression is divided equally into bins so user can view distribution relationship between gene expression and annotations.

The diagram is also shown in an interactive way so that user can change the layout by selecting several items (e.g., thin or thick on color bar, small or large space) from the panel. Also, the user can drag these small boxes on the plot to get preferred layout and save it as high resolution SVG figure.

In addition, when you hover over mouse on a box, you can get detailed information about the source and target of flow

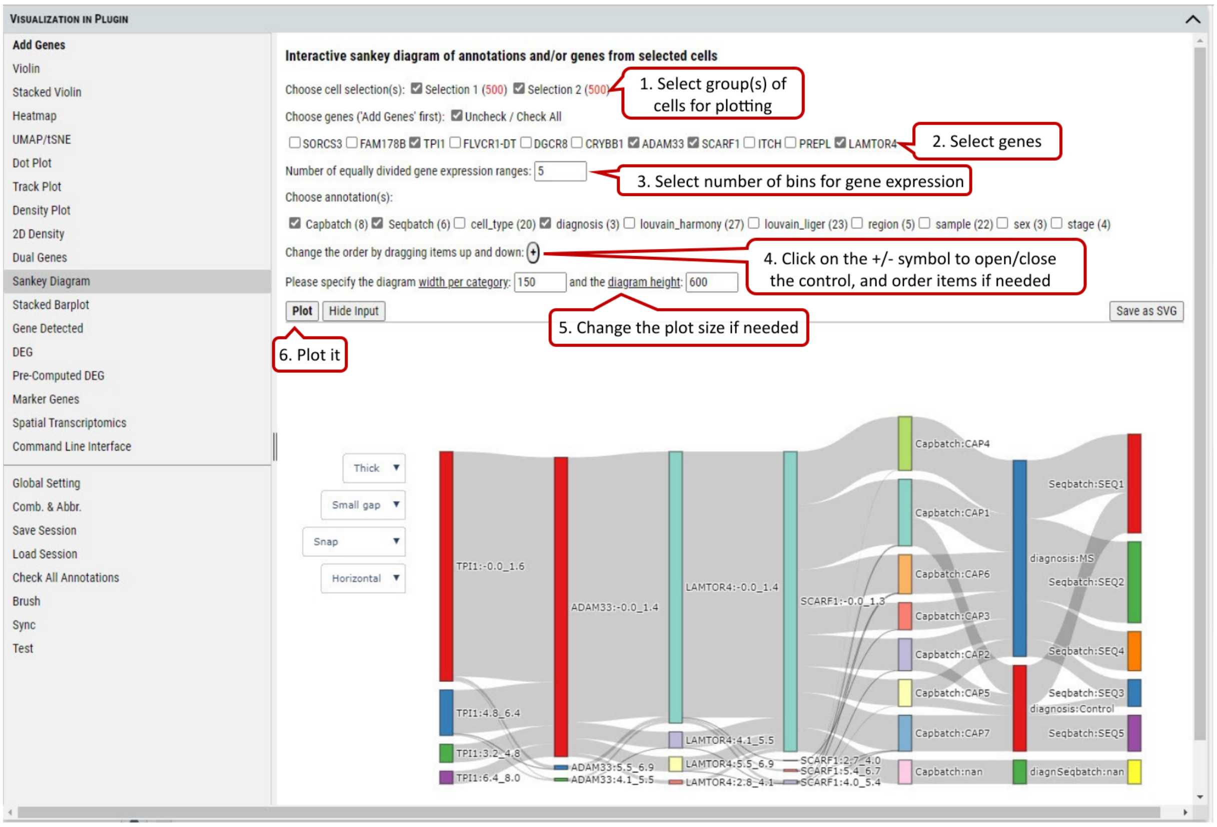

Supplementary Fig. 16. Sankey diagram provides quick and easy way to explore the inter-dependent relationship of variables.

#### 2.17 VIP – Stacked Barplot

To show the distribution of cells among categories of an annotation and/or ranges of expression of agene. Only two factors from annotations or genes can be chosen. The plot allows user to explore the distribution of cells in different views interactively.

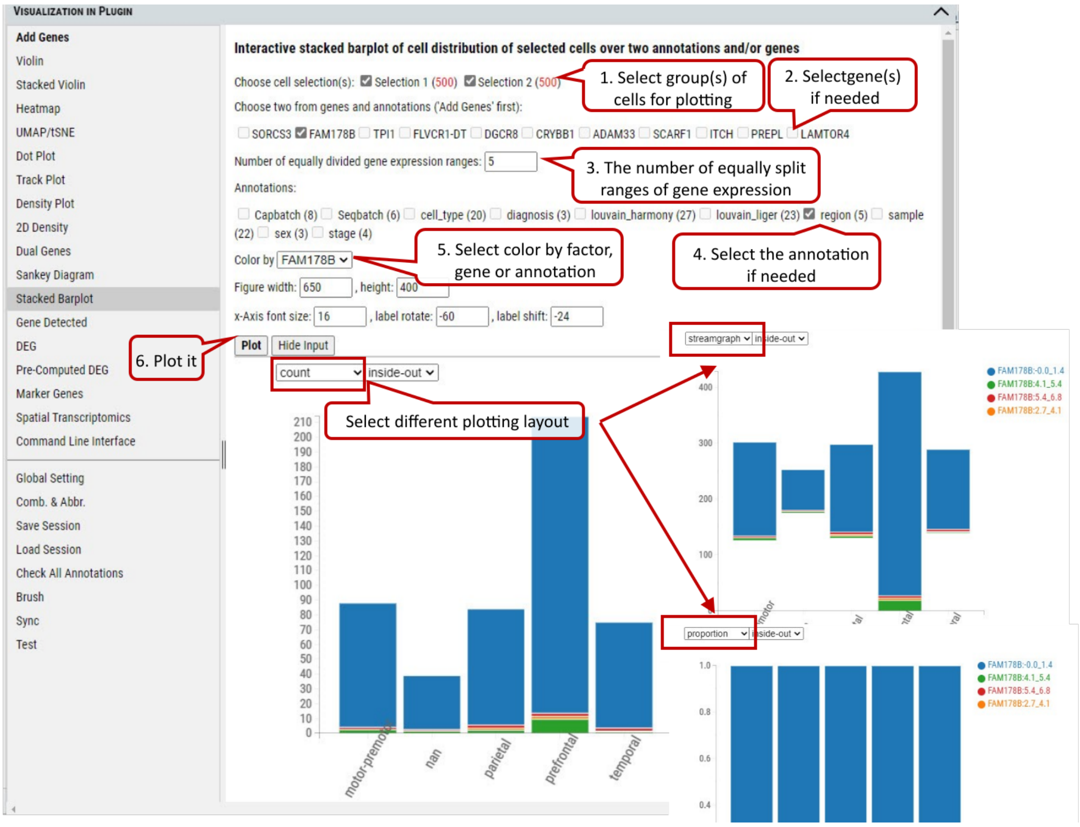

Supplementary Fig. 17. The distribution of cells in the selected group(s) regarding categories of an annotation and expression ranges of a gene by three different layout:count, streamgraph, proportion.

#### 2.18 VIP – Gene Detected

To show the number of genes expressed above the specified expression cut-off in the selected group(s) of cells.

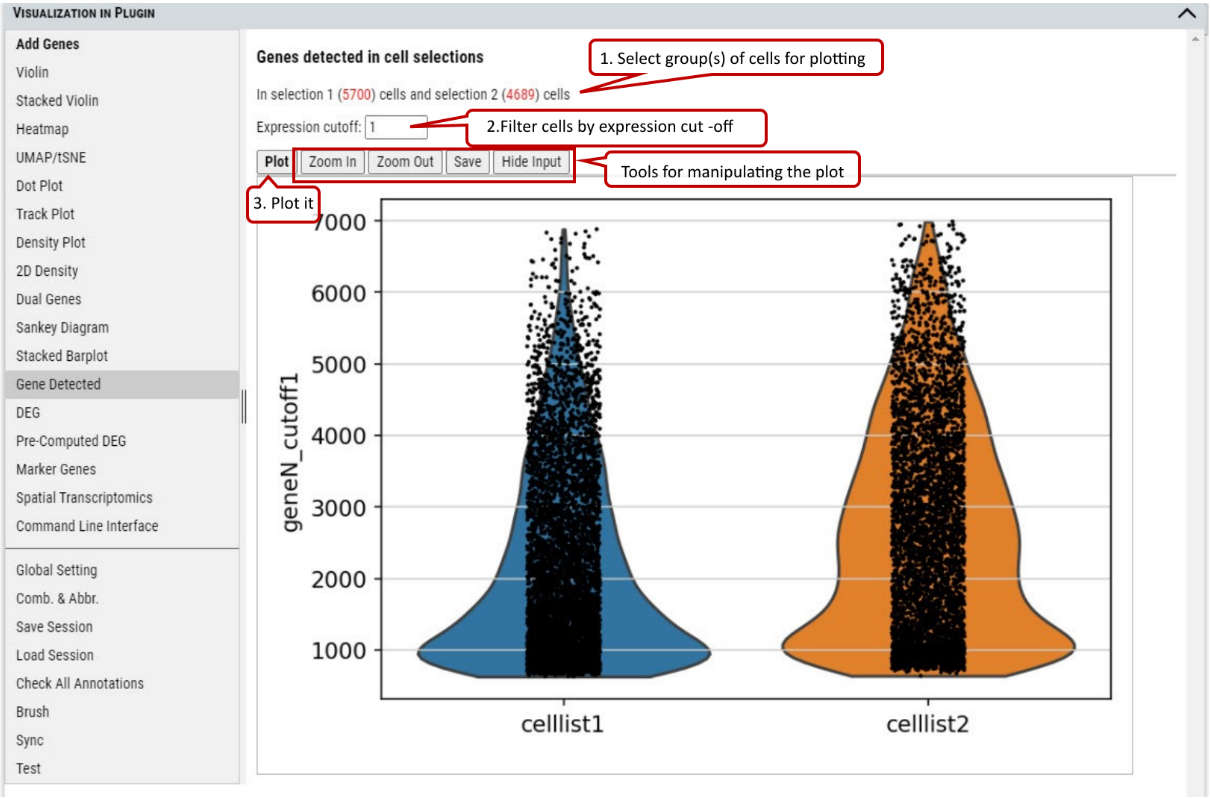

Supplementary Fig. 18. The number of genes with expression over the cut-off in the cells from the selected group(s) of cells.

#### 2.19 VIP – DEG (Differential Expressed Genes)

Besides plotting functions, cellxgene VIP also provides differential analysis between two selected groups of cells to identify differential expressed genes.

Three differential analysis statistical test methods are provided including Welch's t-test, Wilcoxon rank test and Wald's test. The statistical test results are presented in a table format including log2 Fold change, p-value and adjusted p-value (i.e, FDR value). Please note, we provide users with simple test methods for quick exploration within the interactive framework. However, there would be covariates need to be considered in a proper statistical test. Please consult your stats experts for appropriate test by using the right test method and right model.

Volcano plotting is also provided to show the log2FC vs. -log10(FDR) relationship for all genes. The user can select the gene(s) from the pre-selected gene list to be highlighted with text in the volcano plot.

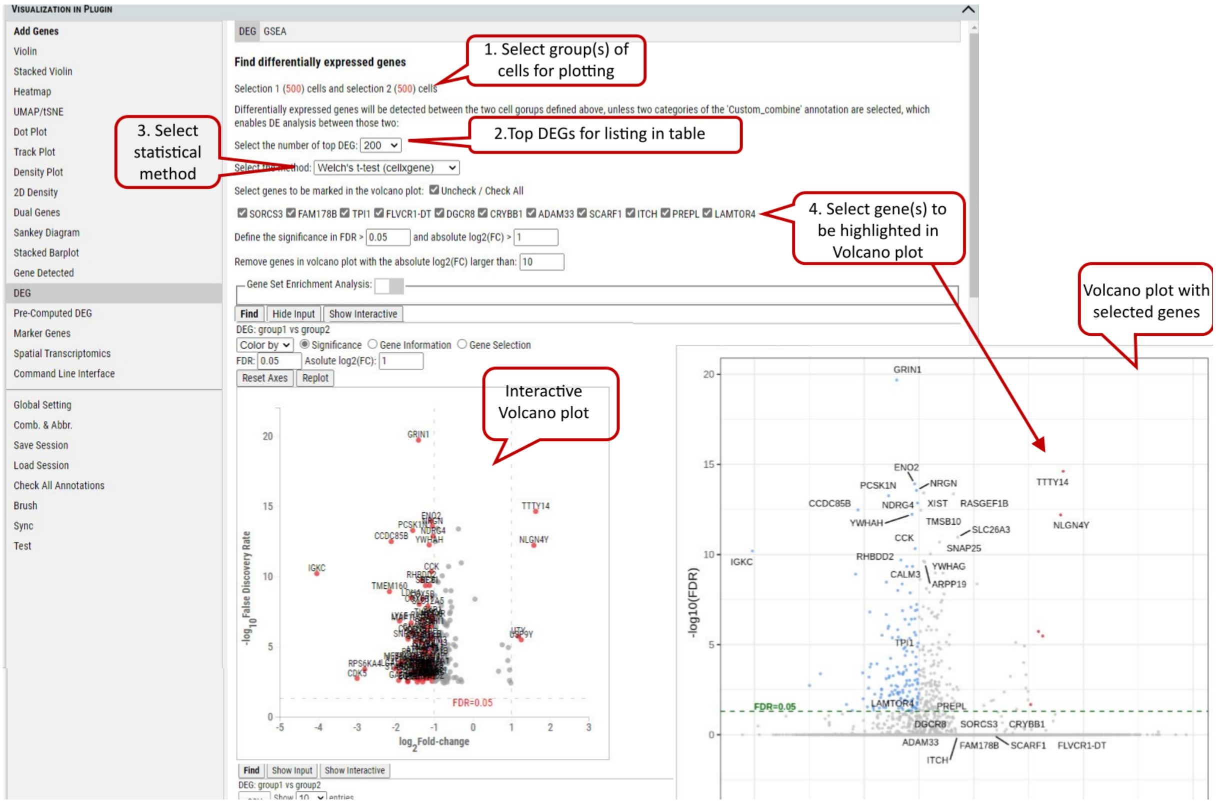

Supplementary Fig. 19. DEG analysis between the selected group(s) with volcano plots

Note: The data used by DEG is unscaled (please refer to description of the dataset to find out what preprocessing was done on the data). Scaling control in the Global Setting does not apply to DEG. The three methods: 'Welch's t-test' uses t-test (assuming underlining data with normal distributions) this uses cellxgene t-test implementation, 'Wilcoxon rank test' uses Wilcoxon rank-sum test (does not assume known distributions, non-parametric test) and 'Wald's test' uses Wald Chi-Squared test which is based on maximum likelihood. 'Wilcoxon rank test' and 'Wald's test' use diffxpy's implementation. Gene set enrichment analysis (GSEA) can be enabled for the DEGs on the gene sets of interest.

#### 2.20 VIP - Pre-computed DEG

In addition, cellxgene VIP shows the differential analysis within some pre-computed annotated groups.

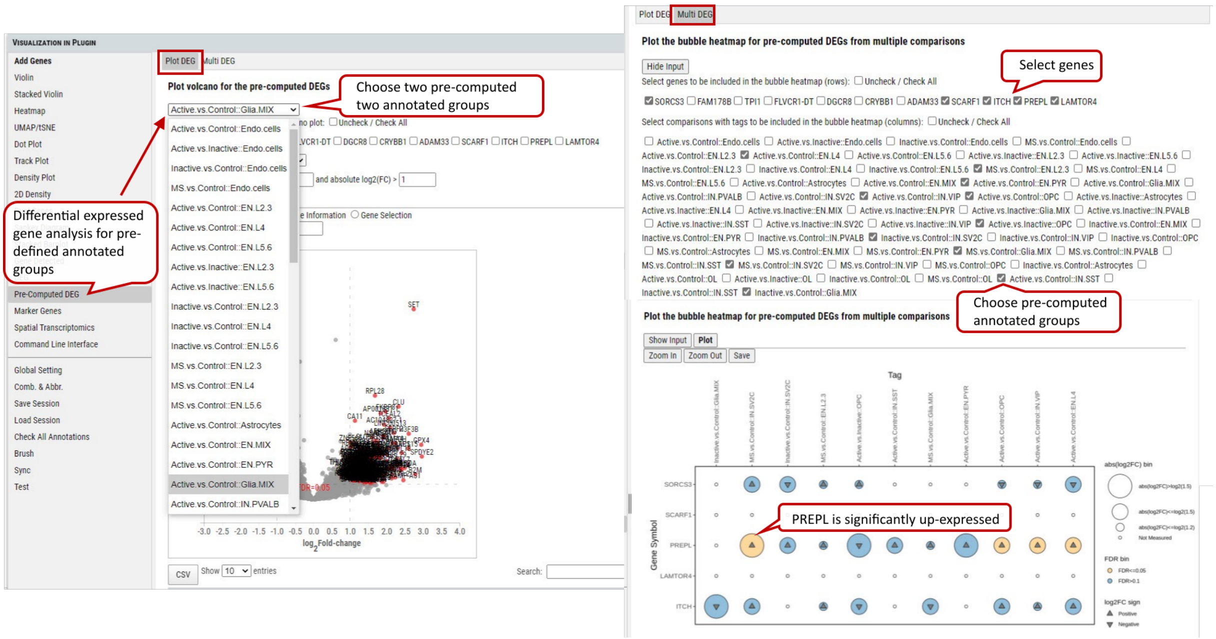

Supplementary Fig. 20. Pre-computed DEG analysis between/ among the selected pre-defubed group(s) with volcano plots and bubble heatmap.

#### 2.21 VIP – Marker Genes

This functional module allows user to identify marker genes in the selected group(s) (more than 2, if 2 groups, please use DEG) of cells by annotation categories.

Four methods are provided for detecting marker genes including logreg, t-test, Wilcoxon, and t-testoverest_var. For each identified marker gene, the gene name, scores (the z-score underlying the computation of a p-value for each gene for each group) and assigned group are listed in the output table.

In each annotation category, top ranked marker genes (this example shows top 2) will be plotted by score in comparison to the rest of the categories.

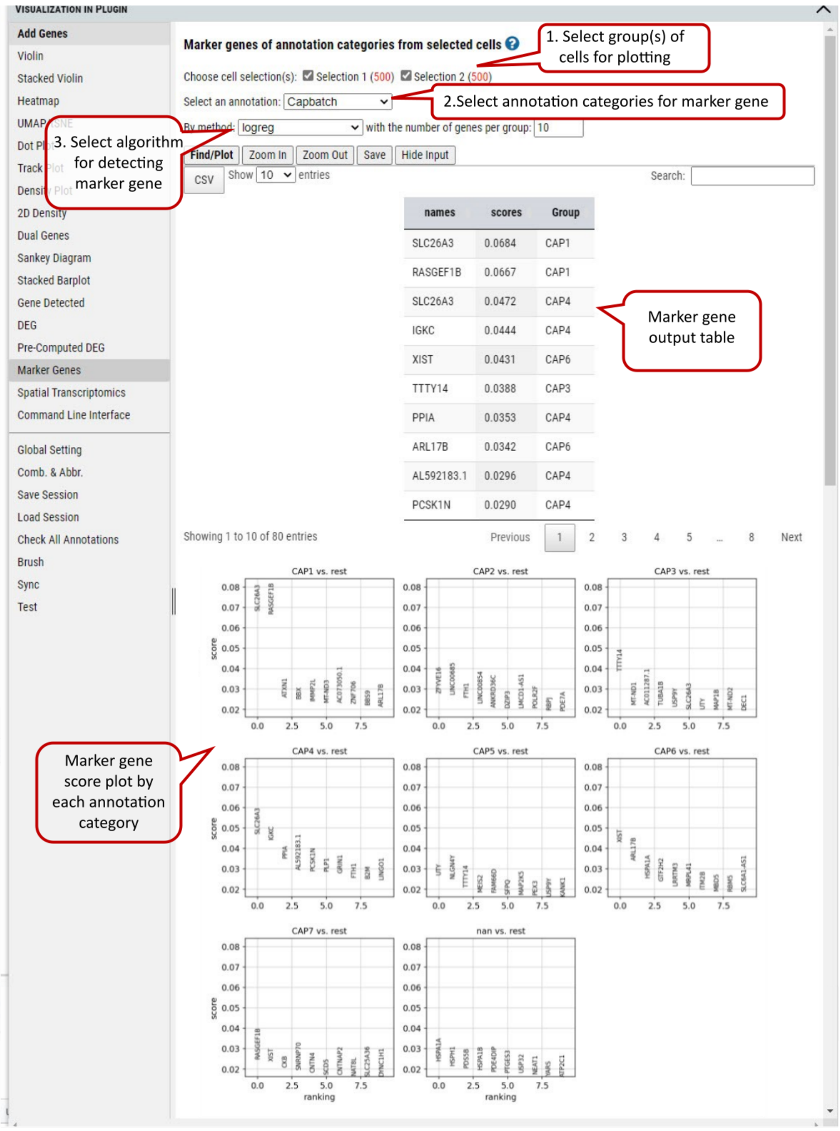

Supplementary Fig. 21. Marker genes detection in the selected group(s) of cells regarding the selected annotation categories.

Note: The four methods implementations by calling scanpy.tl.rank_genes_groups function: 'logreg' uses logistic regression, 't-test' uses t-test, 'wilcoxon' uses Wilcoxon rank-sum, and 't-test_overestim_var' overestimates variance of each group.

#### 2.22 VIP - Spatial Transcriptomics

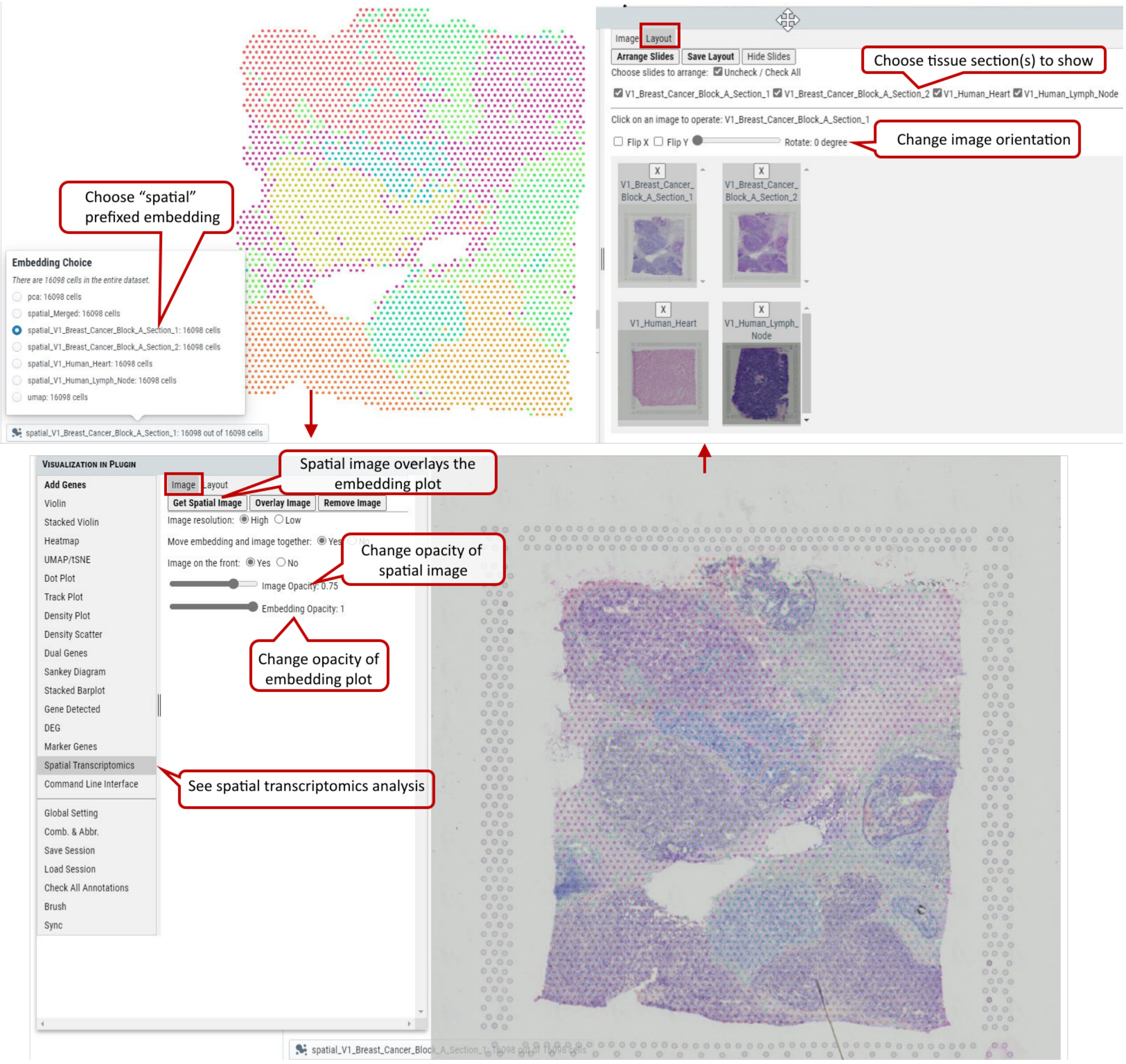

Supplementary Fig. 22. Spatial transcriptomics analysis.

#### 2.23 VIP – Command Line Interface

Although cellxgene VIP provides a rich set of visualization modules as shown above, command line interface is also built to allow unlimited visualization and analytical capabilities by power user who know how to program in Python / R languages.

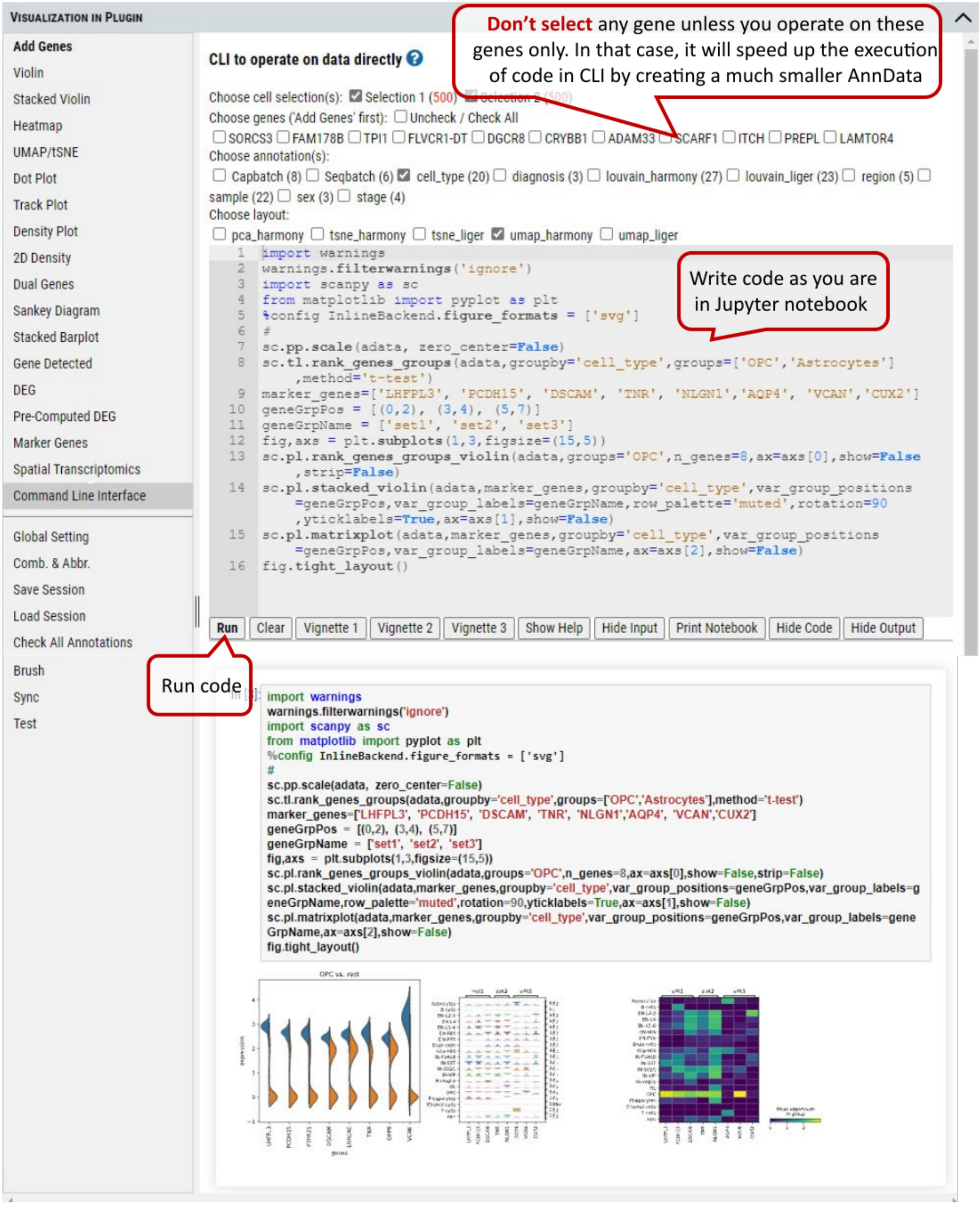

Supplementary Fig. 23. Command line interface for the user to program for advanced plotting and statistical analysis.

Note: In CLI the AnnData (adata) object is available by default, and it is processed as 'Description' of the dataset states (i.e.: normalized and log transformed, but not scaled etc.). Settings in 'Global Setting' tab won't apply to CLI.

#### 2.24 VIP – Comb. & Abbr.

The user can combine multiple annotations to create a combinatorial annotation to group cells in many plotting modules, e.g., stacked violin and dot plot. Firstly, the user clicks 'uncheck all annotations' and, secondly goes to the annotation panel in main window to select annotation categories to be combined, e.g., diagnosis (Control and MS) combined with sex (female and male). After clicking on 'Create' button, all possible combinatorial names will be listed and 'Custom_combine' will be automatically available as an option in 'Group by' drop down menu of many plotting functions.

The user can also rename each annotation by creating abbreviations to shorten axis labels in figures.

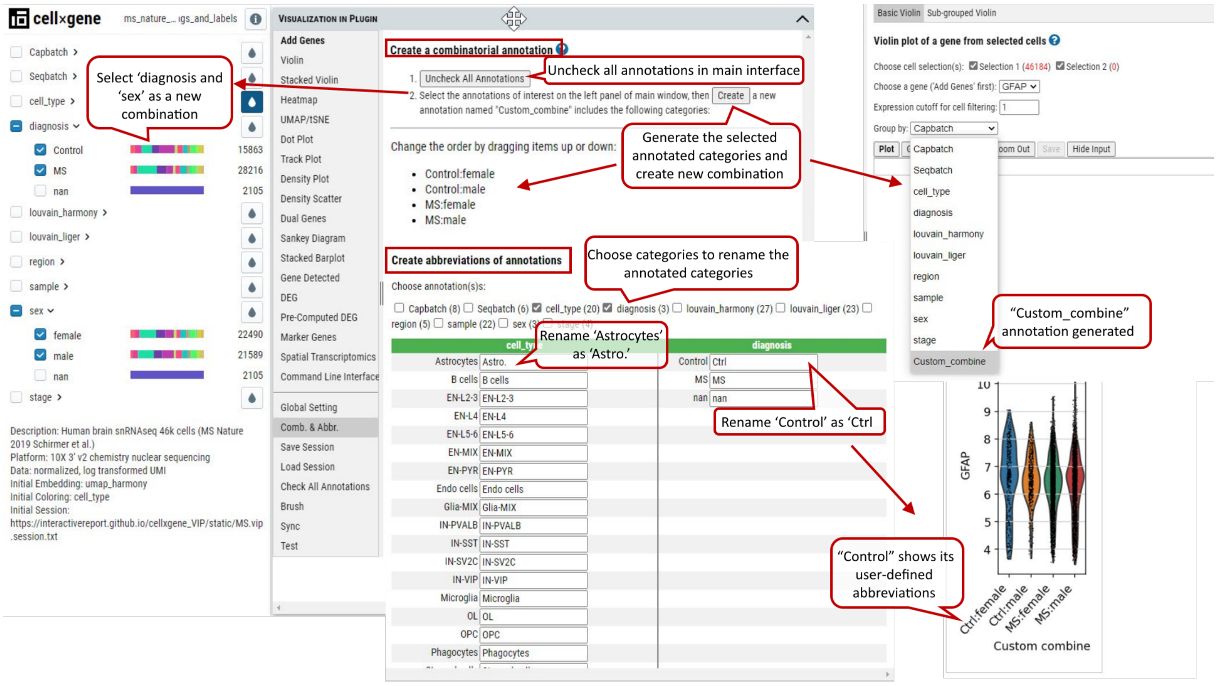

Supplementary Fig. 24. Comb. & Abbr. function allows user to create new annotation by combining multiple annotations and abbreviations to shorten axis labels in figures especially when custom combinatorial names are used.

#### 2.25 VIP – Other Functions

There are other convenient functions available to user, such as 'Save' or 'Load' session, 'Check All Annotations' and 'Brush'.

'Save' or 'Load' session are used to save the current cell selections and parameter settings to text file or load a previously saved session file in the tool for visualization.

'Check All Annotations' is used to check all categorical selection boxes of annotations on the left panel.

'Brush' is to display exactly these selected ranges from histograms of variables on the right panel in a nice table that is not available in original cellxgene.

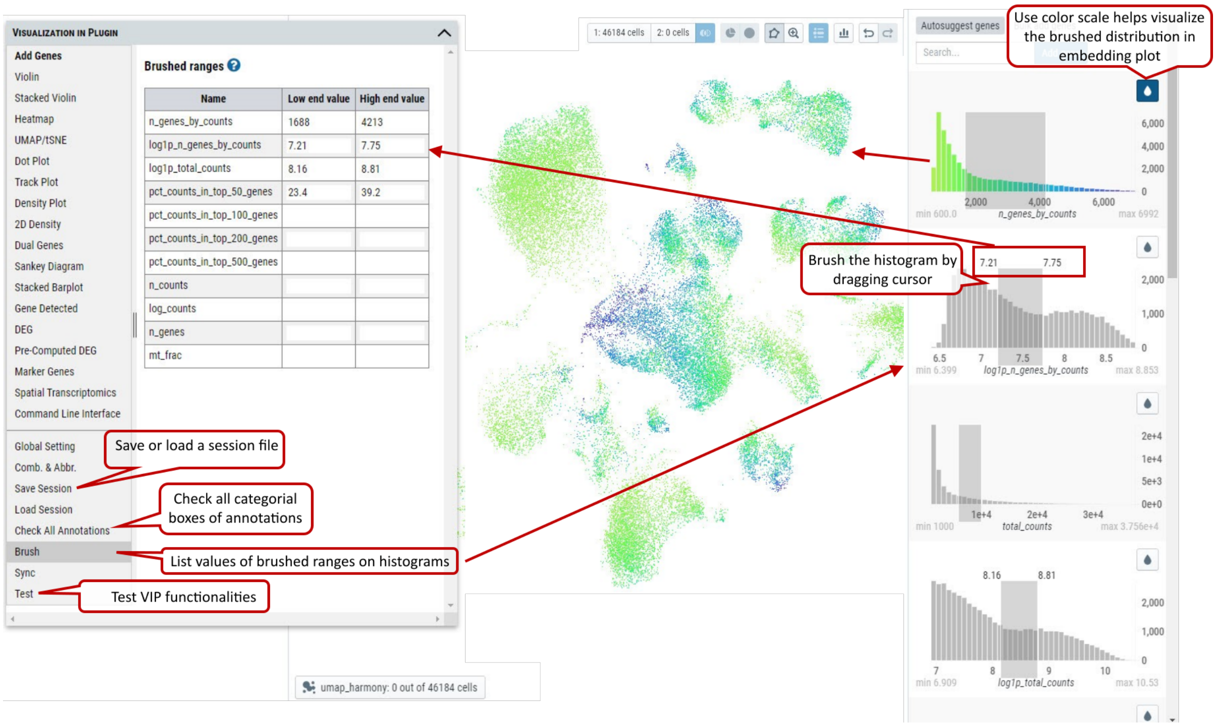

Supplementary Fig. 25. Other functions allow user to 'Save' or 'Load' session information, 'Check All Annotations' and show values of brushed ranges on histograms.

### 3 Methods

#### 3.1 Client-side Integration by a jsPanel Window (VIP)

Related lines in config.sh file.

```
sed -i "s|<div id=\"root\"></div>|$(sed -e 's/[&\\/]/\\&/g; s/|/\\|/g; s/$/\\/;' -e
↪  '$s/\\$//' index_template.insert)\n&|" "cellxgene/client/index_template.html"
```

The content of the index_template.insert file.

```
<script
↪  src="https://interactivereport.github.io/cellxgene_VIP/static/jquery.min.js"></script>
<script src="https://d3js.org/d3.v4.min.js"></script>
<script
↪  src="https://interactivereport.github.io/cellxgene_VIP/static/stackedbar/d3.v3.min.js"></script>
<link
↪  href="https://interactivereport.github.io/cellxgene_VIP/static/jspanel/dist/jspanel.css"
↪  rel="stylesheet">
<script
↪  src="https://interactivereport.github.io/cellxgene_VIP/static/jspanel/dist/jspanel.js"></script>
<script
↪  src="https://interactivereport.github.io/cellxgene_VIP/static/jspanel/dist/extensions/modal/jspa
<script
↪  src="https://interactivereport.github.io/cellxgene_VIP/static/jspanel/dist/extensions/tooltip/js
<script
↪  src="https://interactivereport.github.io/cellxgene_VIP/static/jspanel/dist/extensions/hint/jspan
<script
↪  src="https://interactivereport.github.io/cellxgene_VIP/static/jspanel/dist/extensions/layout/jsp
<script
↪  src="https://interactivereport.github.io/cellxgene_VIP/static/jspanel/dist/extensions/contextmen
<script
↪  src="https://interactivereport.github.io/cellxgene_VIP/static/jspanel/dist/extensions/dock/jspan
```

```
<script>
// execute JavaScript code in panel content
var setInnerHTML = function(elm, html) {
    elm.innerHTML = html;
    Array.from(elm.querySelectorAll('script')).forEach( oldScript => {
        const newScript = document.createElement('script');
        Array.from(oldScript.attributes)
        .forEach( attr => newScript.setAttribute(attr.name, attr.value) );
        newScript.appendChild(document.createTextNode(oldScript.innerHTML));
        oldScript.parentNode.replaceChild(newScript, oldScript);
    });
}
var plotPanel = jsPanel.create({
    panelSize: '190 0',
    position: 'left-top 160 6',
    dragit: { containment: [-10, -2000, -4000, -2000] }, // set dragging range of VIP
    ↪ window
    boxShadow: 1,
    border: "solid #D4DBDE thin",
    contentOverflow: 'scroll scroll', // adding scrolling bars
    headerControls:{
      close: 'remove',
      minimize: 'remove',
      maximize: 'remove'
    },
    headerTitle: function () {return '<strong>Visualization in Plugin</strong>'},
    contentAjax: {
        url: window.location.href.replace(/\\\/+$/,'')+'/static/interface.html',
        done: function (panel) {
               setInnerHTML(panel.content, this.responseText);
        }
    },
    onwindowresize: function(event, panel) {
        var jptop = parseInt(this.currentData.top);
        var jpleft = parseInt(this.currentData.left);
        if (jptop<-10 || window.innerHeight-jptop<10 || window.innerWidth-jpleft<10 ||
        ↪ jpleft+parseInt(this.currentData.width)<10) {
            this.reposition("left-top 160 6");
        }
    },
    onunsmallified: function (panel, status) {
        this.reposition('center-top -370 180');
        this.resize({ width: 740, height: function() { return Math.min(480,
        ↪ window.innerHeight*0.6);} });
    },
    onsmallified: function (panel, status) {
        this.reposition('left-top 160 6');
        this.style.width = '190px';
    }
}).smallify();
plotPanel.headerbar.style.background = "#D4DBDE";
</script>
```

All functional VIP HTML and JavaScript code will be in a new file called “interface.html”, which is out of cellxgene code base.

#### 3.2 Server-side Integration

Related in config.sh file.

```
echo '
from server.app.VIPInterface import route
@webbp.route("/VIP", methods=["POST"])
def VIP():
    return route(request.data,current_app.app_config)' >> cellxgene/server/app/app.py

cd cellxgene
make pydist
make install-dist
cd ..

## finished setting up ------
./update.VIPInterface.sh all
```

The content of update.VIPInterface.sh file.

```
#!/usr/bin/env bash
if [ -n "$1" ]; then
echo "usually update once"
fi

## finished setting up ------
strPath="$(python -c 'import site; print(site.getsitepackages()[0])')"
strweb="${strPath}/server/common/web/static/."

cp VIPInterface.py $strPath/server/app/.
cp interface.html $strweb
cp vip.env $strPath/server/app/. 2>/dev/null | true

cp fgsea.R $strPath/server/app/.
mkdir -p $strPath/server/app/gsea
cp gsea/*gmt $strPath/server/app/gsea

if [ -n "$1" ]; then
  cp Density2D.R $strPath/server/app/.
  cp bubbleMap.R $strPath/server/app/.
  cp violin.R $strPath/server/app/.
  cp volcano.R $strPath/server/app/.
  cp browserPlot.R $strPath/server/app/.
  if [ "$(uname -s)" = "Darwin" ]; then
    sed -i .bak "s|route(request.data,current_app.app_config,
    ↪  \"/tmp\")|route(request.data,current_app.app_config)|"
    ↪  "$strPath/server/app/app.py"
    sed -i .bak "s|MAX_LAYOUTS *= *[0-9]\+|MAX_LAYOUTS = 300|"
    ↪  "$strPath/server/common/constants.py"
  else
    sed -i "s|route(request.data,current_app.app_config,
    ↪  \"/tmp\")|route(request.data,current_app.app_config)|"
    ↪  "$strPath/server/app/app.py"
    sed -i "s|MAX_LAYOUTS *= *[0-9]\+|MAX_LAYOUTS = 300|"
    ↪  "$strPath/server/common/constants.py"
  fi

  find ./cellxgene/server/ -name "decode_fbs.py" -exec cp {} $strPath/server/app/. \;
fi
```

```
echo -e "\nls -l $strweb\n"
ls -l $strweb
```

##### 3.3 Communication between VIP and cellxgene web GUI

Cellxgene client utilizes React Redux that is the official React binding for Redux. It lets your React components read data from a Redux store, and dispatch actions to the store to update data.

So, this following code in config.sh appends "window.store = store;" to the end of client/src/reducers/index.js of cellxgene source code to expose the store to the browser.

```
echo -e "\nwindow.store = store;" >> cellxgene/client/src/reducers/index.js
```

By doing this, Redux store holding client data and user selections are visible to VIP to access variables and dispatch actions to control cellxgene user interface. For example,

- Unselect / select a feature. GUI is refreshed automatically after dispatching.

```
window.store.dispatch({type: "categorical metadata filter deselect", metadataField:
↪  "louvain", categoryIndex: 5})
window.store.dispatch({type: "categorical metadata filter select", metadataField:
↪  "louvain", categoryIndex: 5})
```

- Get the state of a just finished action and synchronize gene input and cell selections from main window to VIP if corresponding action was performed.

```
const unsubscribe = window.store.subscribe(() => {
  if (window.store.getState()["@@undoable/filterState"].prevAction) {
    actionType = window.store.getState()["@@undoable/filterState"].prevAction.type;
    if (actionType.includes("user defined gene success") ||
    actionType.includes("store current cell selection as differential set")) {
      sync();
      }
  }
});
```

##### 3.4 Diffxpy Integration

This is the sample pseudocode, please see VIPInterface.py for actual implementation.

```
import scanpy as sc
import pandas as pd
import diffxpy.api as app
# set 1 of cells as cell1; set 2 of cells as cell2

with app.get_data_adaptor() as data_adaptor:
  X1 = data_adaptor.data.X[cell1]
  X2 = data_adaptor.data.X[cell2]

adata = sc.AnnData(pd.concat([X1,X2]),pd.DataFrame(['grp1' for i in
↪  range(X1.shape[0])]+['grp2' for i in range(X2.shape[0])],columns=['comGrp']))
deg = de.test.two_sample(adata,'comGrp').summary()
#deg is a dataframe contains the folloing columns ['gene','log2fc','pval','qval']
```

#### 3.5 Create a h5ad file from Seurat object

First, export the following from Seurat object in R: **expression matrix (assume normalized), metadata and coordinates (pca, tsne, umap) as separate txt files.**

Next in Python, create an AnnData object from 10x (scanpy.read_h5ad function) as a starting point. Then replace the expression matrix, meta data and coordinates as shown in the following Python code block to generate a h5ad file.

```
import sys
import scanpy as sc
import pandas as pd
import numpy as np
import seaborn as sns
from numpy import ndarray, unique
from scipy.sparse.csc import csc_matrix

adata= sc.read_h5ad("previous generated .h5ad")

# read clustering res
xpca = pd.read_csv("./data/harmony_clustered.h5ad.pca_coordinates.txt", sep='\t',
↪  encoding='utf-8')
xtsne = pd.read_csv("./data/harmony_clustered.h5ad.tsne_coordinates.txt", sep='\t',
↪  encoding='utf-8')
xumap = pd.read_csv("./data/harmony_clustered.h5ad.umap_coordinates.txt", sep='\t',
↪  encoding='utf-8')
xobs = pd.read_csv("./data/harmony_clustered.h5ad.meta_data.txt", sep='\t',
↪  encoding='utf-8')

xpca.set_index('index', inplace=True)
xtsne.set_index('index', inplace=True)
xumap.set_index('index', inplace=True)
xobs.set_index('index', inplace=True)

adata.obsm['X_pca'] = np.array(xpca.loc[adataRaw.obs.index])

adata.obsm['X_tsne'] = np.array(xtsne.loc[adataRaw.obs.index])
adata.obsm['X_umap'] = np.array(xumap.loc[adataRaw.obs.index])
adata.obs = xobs.loc[adataRaw.obs.index] # this is a pandas dataframe

# read in expression matrix as numpy.ndarray
exp_mat = np.loadtxt(fname ="expression matrix .txt")
adata.X = exp_mat

# convert dense matrix into sparse matrix to save storage space and memory usage
adata.X = csc_matrix(adata.X)_matrix

# add short description and initial graph settings. "|" and "by" are delimiters for
↪  VIP to parse the initial settings. Please follow the same rule for your own h5ad
↪  files.
adata.obs['>Description'] = ['Human brain snRNAseq 46k cells (MS Nature 2019 Schirmer
↪  et al.); data normalized, log transformed and scaled UMI; platform - 10X v2
↪  chemistry | embedding by umap; color by cell_type']*adata.n_obs

# Then last step to save h5ad:
adata.write_h5ad("final output.h5ad")
```

When the h5ad file is uploaded to cellxgeneVIP, AnnData.X matrix is to be used for visualization and DEG analysis. By default, the data (e.g, raw count matrix) is assumed to be unscaled , however, if the data have been scaled or normalized, the user needs to turn off the option 'Scale data to unit variance for plotting:' in 'Global Setting'.

### 4 Helpful Tips

#### 4.1 Handle nulls in categorical annotation

Such nulls in categorical annotation would cause trouble in VIP because it cannot be converted to string. Here is how to handle it, let's call the annotation X_annotation:

```
# Cast to str from categorical
adata.obs = adata.obs.astype({'X_annotation':'str'})

# replace all of nan by 'nan'
adata.obs["X_annotation "][adata.obs["X_annotation "].isnull()] = 'nan'
```

#### 4.2 Display full traceback stack for debugging in VIP

It follows the global setting. Please set "—verbose" to launch cellxgene server.

#### 4.3 Pitfall of using special characters

In the mode which allows user to create manual annotation in cellxgene, user should try to avoid using hyphen ("-") in name label. It would cause client-side issue. Please try to use underscores.

#### 4.4 Potential use for bulk or pseudo bulk sample dataset

Once the data matrix is replaced by sample x gene matrix, cellxgene VIP framework can handle regular bulk / pseudobulk RNAseq datasets. Simply replace "cells" by "samples". All plotting functions sould still work.

#### 4.5 Common mistakes in naming

- "B internediate" with extra space in the end of a cell type
- "CD4 naive" where "n" should be "N" to match the cell type used in the h5ad file

#### 4.6 Loading h5ad file generated by SeuratDisk from R

SeuratDisk provides a convenient way to convert a Seurat object into the h5ad format. However, there are cases cellxgene VIP window would be blank when loading such h5ad file. To fix the issue, simply run the following python code to re-generate the h5ad file, which should solve the blank VIP window issue.

```
import scanpy as sc
adata = sc.read_h5ad(<input h5ad>)
adata.__dict__['_raw'].__dict__['_var'] =
↪  adata.__dict__['_raw'].__dict__['_var'].rename(columns={'_index': 'features'})
```

### 5 Web Resource

#### 5.1 Cellxgene VIP

- **cellxgene VIP source code**: https://github.com/interactivereport/cellxgene_VIP
- **cellxgene VIP demo sites**: https://cellxgenevip-ms.bxgenomics.com [Schirmer / Rowitch MS, Human brain snRNAseq 46k cells, Nature 2019 Schirmer et al]

#### 5.2 Cellxgene

- **cellxgene**: https://github.com/chanzuckerberg/cellxgene
- **cellxgene tutorial**: https://cellgeni.readthedocs.io/en/latest/visualisations.html
- **cellxgene features**: https://chanzuckerberg.github.io/cellxgene/posts/gallery

- **cellxgene data preparation**: https://chanzuckerberg.github.io/cellxgene/posts/prepare.html

#### 5.3 Python packages

- **scanpy**: https://github.com/theislab/scanpy
- **scanpy plots**: https://scanpy-tutorials.readthedocs.io/en/latest/visualizing-marker-genes.html
- **diffxpy**: https://github.com/theislab/diffxpy
- **diffxpy tests**: https://diffxpy.readthedocs.io/en/latest/tutorials.html#differential-testing

#### 5.4 Others

- **Benchmarking of interactive data visualization of single-cell RNAseq data — Batuhan Cakir** : https://www.youtube.com/watch?v=3nH2xi_Ni6I
- **sceasy**: convertor of scRNA-Seq data formats https://github.com/cellgeni/sceasy
- **jupytext**: https://jupytext.readthedocs.io , Jupyter Notebooks as Markdown Documents, Julia, Python or R Scripts
- **nbconvert**: https://nbconvert.readthedocs.io , Convert a Jupyter .ipynb notebook document file into another static format including HTML, LaTeX, PDF, Markdown, and more
- **R magic**: https://rpy2.github.io/doc/latest/html/interactive.html#rmagic , Magic command interface for interactive work with R in ipython. %R and %%R are for the line mode where one line of R code will be executed, and the cell mode where a block of R code will run, respectively.
